# Long-term Hematopoietic Transfer of the Anti-Cancer and Lifespan-Extending Capabilities of A Genetically Engineered Blood System by Transplantation of Bone Marrow Mononuclear Cells

**DOI:** 10.1101/2023.04.21.537849

**Authors:** Jing-Ping Wang, Chun-Hao Hung, Yao-Huei Liou, Ching-Chen Liu, Kun-Hai Yeh, Keh-Yang Wang, Zheng-Sheng Lai, Biswanath Chatterjee, Tzu-Chi Hsu, Tung-Liang Lee, Yu-Chiau Shyu, Pei-Wen Hsiao, Liuh-Yow Chen, Trees-Juen Chuang, Chen-Hsin Albert Yu, Nah-Shih Liao, Che-Kun James Shen

**Author notes:** Corresponding Authors, Please send all correspondence to: Che-Kun James Shen The PhD Program for Neural Regenerative Medicine Taipei Medical University Taipei 115, Taiwan, ROC Yu-Chiau Shyu Community Medicine Research Center, Chang Gung Memorial Hospital, Keelung Branch, Keelung 204, Taiwan (R.O.C.) Chun-Hao Hung The PhD Program for Neural Regenerative Medicine Taipei Medical University Taipei 115, Taiwan, ROC. Contributed equally (Jing-Ping Wang, Chun-Hao Hung, Trees-Juen Chuang).

## Abstract

A causal relationship exists among the aging process, organ decay and dis-function, and the occurrence of various diseases including cancer. A genetically engineered mouse model, termed *Klf1*^K74R/K74R^ or *Klf1*(K74R), carrying mutation on the well-conserved sumoylation site of the hematopoietic transcription factor KLF1/ EKLF has been generated that possesses extended lifespan and healthy characteristics including cancer resistance. We show that the healthy longevity characteristics of the *Klf1*(K74R) mice, as exemplified by their higher anti- cancer capability, are likely gender-, age- and genetic background-independent. Significantly, the anti-cancer capability, in particular that against melanoma as well as hepatocellular carcinoma, and lifespan-extending property of *Klf1*(K74R) mice could be transferred to wild-type mice via transplantation of their bone marrow mononuclear cells at young age of the latter. Furthermore, NK(K74R) cells carry higher *in vitro* cancer cell-killing ability than wild type NK cells. Targeted/global gene expression profiling analysis has identified changes of the expression of specific proteins, including the immune checkpoint factors PDCD and CD274, and cellular pathways in the leukocytes of the *Klf1*(K74R) that are in the directions of anti-cancer and/or anti-aging. This study demonstrates the feasibility of developing a transferable hematopoietic/ blood system for long-term anti-cancer and, potentially, for anti-aging.

## Introduction

Aging of animals, including humans, is accompanied by lifespan-dependent organ deterioration and the occurrence of chronic diseases such as cancer, diabetes, cardiovascular failure and neurodegeneration^1,2^. To extend healthspan and lifespan, various biomedical- and biotechnology-related strategies have been intensively developed and applied, including the therapy of different diseases such as cancer^3–5^. The hematopoietic/ blood system is an important biomedical target for anti-aging and anti-cancer research development. Multiple blood cell lineages arise from hematopoietic stem cells (HSCs), with the lymphoid lineage giving rise to T, B, and natural killer (NK) cell populations, whereas the myeloid lineage differentiates into megakaryocytes, erythrocytes, granulocytes, monocytes and macrophages^6–8^. The genetic constituents and homeostasis of the hematopoietic system are regulated epigenetically and via environmental factors to maintain animal health^6,9^.

EKLF, also named KLF1, is a Krüppel-like factor that is expressed in a range of blood cells including erythrocytes, megakaryocytes, T cells, NK cells, as well as in various hematopoietic progenitors including common-myeloid-progenitor (CMP), megakaryocyte-erythroid-progenitor (MEP), and granulocyte-macrophage-progenitor (GMP)^10–12^. The factor regulates erythropoiesis^13^ and the differentiation of MEP to megakaryocytes and erythrocytes^10,14^ as well as of monocytes to macrophages^15^. KLF1 is also expressed in HSC and regulates their differentiation^16^. The factor can positively or negatively regulate transcription, including the adult β globin genes, through binding of its canonical zinc finger domain to the CACCC motif of the regulatory regions of a diverse array of genes^16–19^. In fact, the zinc finger of KLF1 could recognize/bind to genomic sites with the consensus sequence CCNCNCCC, and recruit different co-activator or co-repressor chromatin remodeling complexes^14,17^.

KLF1 could be sumoylated *in vitro* and *in vivo*, and sumoylation of the lysine at codon 74 of mouse KLF1 altered the transcriptional regulatory function^10^ as well as nuclear import^20^ of the factor. Surprisingly, homozygosity of a single amino acid substitution, lysine(K) to arginine(R), at the sumoylation site of KLF1 results in the generation of a novel mouse model with healthy longevity. These mice, termed *Klf1*^K74R/K74R^ or *Klf1*(K74R), exhibited extended healthspan and lifespan. In particular, the *Klf1*(K74R) mice showed delay of the age-dependent decline of physical performance, such as the motor function and spatial learning/memory capability, and deterioration of the structure/function of tissues including the heart, liver, and kidney. Furthermore, the *Klf1*(K74R) mice appeared to have significantly higher anti-cancer capability than the WT mice^21^.

As described in the following, we have since characterized the high anti-cancer capability of the *Klf1*(K74R) mice with respect to its dependence on the age, gender and genetic background. More importantly, we have demonstrated that the high anti-cancer ability of these genetically engineered mice could be transferred to wild type mice (WT) through hematopoietic transplantation of the bone marrow mononuclear cells (BMMNC). Furthermore, we show that the higher anti-cancer capability and extended life span of *Klf1*(K74R) mice are associated with changes of the global protein expression profile and specific aging-/cancer-associated cellular signaling pathways in their white blood cells (WBC) or leukocytes.

## Result

### Characterization of the cancer resistance of *Klf1*(K74R) mice in relation to age, gender, and genetic background

The *Klf1*(K74R) mice appeared to be cancer resistant to carcinogenesis as manifested by their lower spontaneous cancer incidence (12.5%) in life than WT mice (75%). The *Klf1*(K74R) mutation also protected the mice from metastasis of melanoma cells and it reduced melanoma growth in the subcutaneous cancer cell inoculation assay^21^. Here we used the pulmonary melanoma foci assay, which has been commonly used for quick analysis of early metastasis^22^, to further characterize the higher cancer resistance of the *Klf1*(K74R) mice with respect to the effects of gender/age/genetic background of the mice and the requirement for the homozygosity of the K74R mutation. The cultured melanoma B16-F10 cells used in this assay are poorly immunogenic and highly aggressive thus being useful for monitoring early metastasis in various immunotherapy studies^22,23^.

It appeared that male as well as female *Klf1*(K74R) mice in the B6 genetic background had significantly fewer pulmonary melanoma foci than the corresponding WT mice (Figure 1). Because of this result, we used male mice for all of the studies describe below. First, both young (2 month-old) and aged (24 month-old) *Klf1*(K74R) mice had higher anti-metastasis ability against the injected melanoma cells than WT mice of age-dependent groups (Figure 1A and 1B). Secondly, homozygosity of the K74R substitution was required for the higher cancer resistance of the *Klf1*(K74R) mice (Figure 1-figure supplement 1). Importantly, the *Klf1*(K74R) mice in the FVB background also exhibited high cancer resistance than FVB WT mice by this assay (Figure 1C), suggesting that cancer resistance of *Klf1*(K74R) mice conferred by the homozygous K74R substitution was likely genetic background-independent. Finally, the higher anti-cancer capability of the *Klf1*(K74R) mice did not appear to depend on the arginine at codon 74, since *Klf1*(K74A) mice carrying KèA amino acid substitution at the K74 sumoylation site also exhibited higher anti-metastasis capability than WT mice in the pulmonary foci assay (Figure 1D).

**Figure 1.**
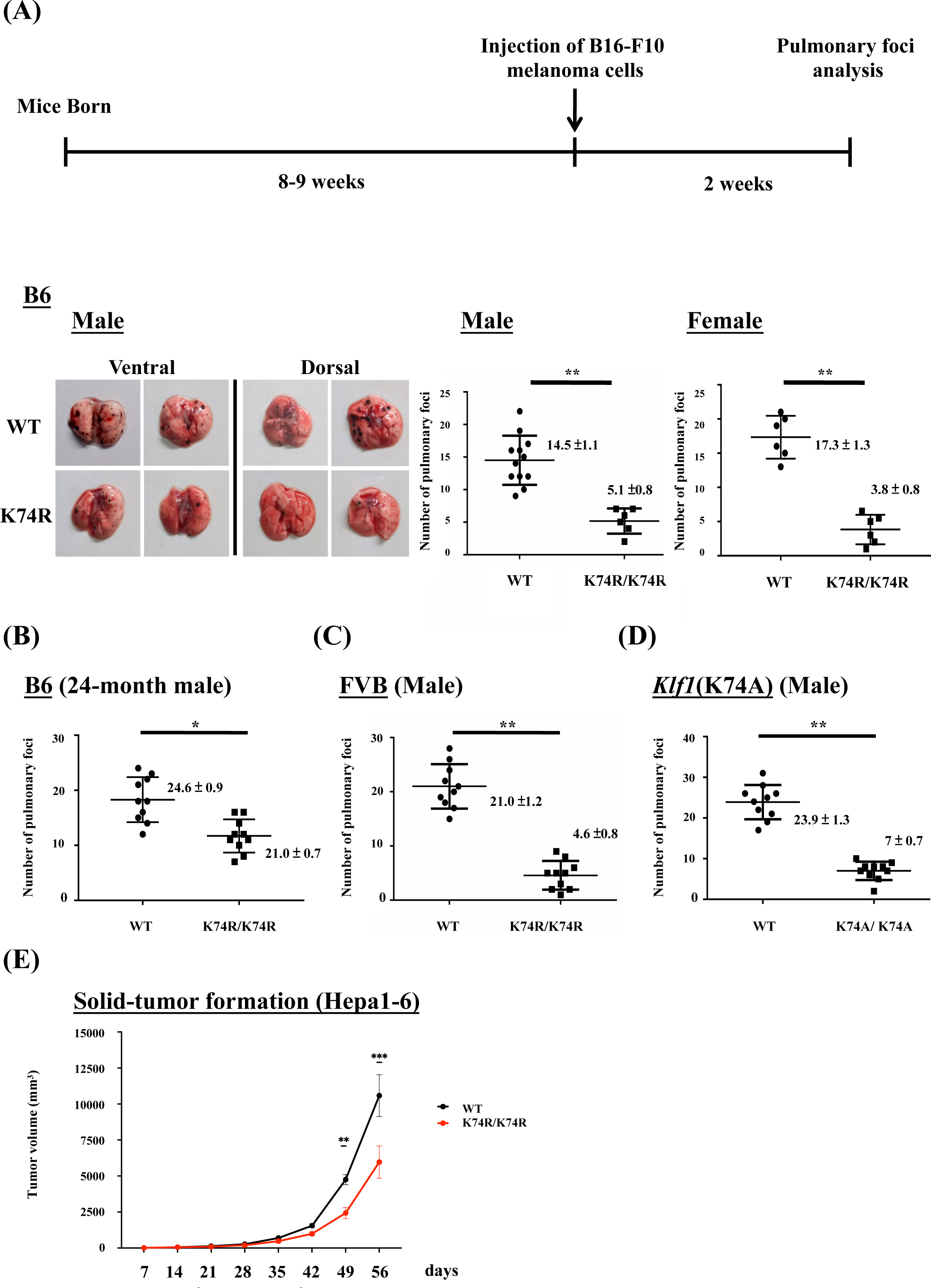
Anti-cancer capability of *Klf1*(K74R) mice as analyzed by the experimental melanoma metastasis assay. (A) Flow chart illustrating the strategy of the pulmonary tumor foci assay. Left panels, representative photographs of pulmonary metastatic foci on the lungs of WT and *Klf1*(K74R) male mice in the B6 background two weeks after intravenous injection of B16-F10 cells (10^5^ cells/ mouse). Statistical comparison of the numbers of pulmonary foci is shown in the two histograms on the right. N=10 (male) and N=7 (female), **, p<0.01. Note that only the numbers of large pulmonary foci (>1mm diameter) were scored. N>6, **, p<0.01. (B) Pulmonary tumor foci assay of 24 month-old WT and *Klf1*(K74R) male mice. Statistical comparison is shown in the two histograms. N=10 (male), *, p<0.05. (C) Pulmonary tumor foci assay of male mice in the FVB background. Statistical comparison is shown in the histograph on the right. N=10, **, p<0.01. (D) Pulmonary tumor foci assay of *Klf1*(K74A) male mice. Statistical comparison of the 3 month-old WT and *Klf1*(K74A) mice numbers of pulmonary foci is shown in the two histograms. N=10 (male), **, p<0.01. (E) The *Klf1*(K74R) mice and WT mice were subcutaneously injected with Hepa1-6 cells to form tumors. The tumor volumes were measured once a week using the formula: length x weight x height x π/6. The curves of tumor growth started to show difference between the *Klf1*(K74R) and WT mice at 49 days after Hepa1-6 cell injection. N>6, *p < 0.05, **p < 0.01 and ***p < 0.001.

The anti-cancer capability of *Klf1*(K74R) mice was not restricted to melanoma. As shown by the subcutaneous inoculation assay of Hepa1-6 cells (Figure 1E), *Klf1*(K74R) mice also carry a higher anti-cancer capability against hepatocellular carcinoma than the WT mice. It thus appears that the *Klf1*(K74R) mice are resistant to the carcinogenesis of a range of different cancers.

### Transfer of the anti-cancer capability and extended lifespan of *Klf1*(K74R) mice to WT mice via BMT

Since KLF1 is a hematopoietic transcription factor expressed not only in mature blood cells but also in HSCs and hematopoietic stem progenitor cells, this cancer resistance may be transferable by means of BMT. This possibility was tested with uses of male mice and a standard BMT protocol^24^. BMMNC were purified from the bone marrow of 2 month-old CD45.2 *Klf1*(K74R) or WT mice and injected into the tail vein of CD45.1 WT recipient mice. Blood replacement of recipient mice with 10Gy γ-irradiation by that of the donor mice reached 90% at 7^th^-week (Figure 2A and 2B). After 2 weeks, the recipient mice were injected with B16-F10 cells and then sacrificed a further 2 weeks later to quantify pulmonary tumor foci. We found that WT mice transplanted with WT BMMNC had similarly high numbers of tumor foci relative to WT mice without BMT (Figure 2C and 1A). However, similar to *Klf1*(K74R) mice challenged with B16-F10 cells (Figure 1B), WT mice that received BMMNC from *Klf1*(K74R) mice presented significantly fewer tumor foci on their lungs (Figure 2C). Notably, BMT using 24 month-old donor mice gave similar result although the effect is relatively small (Figure 2-figure supplement 1A).

**Figure 2.**
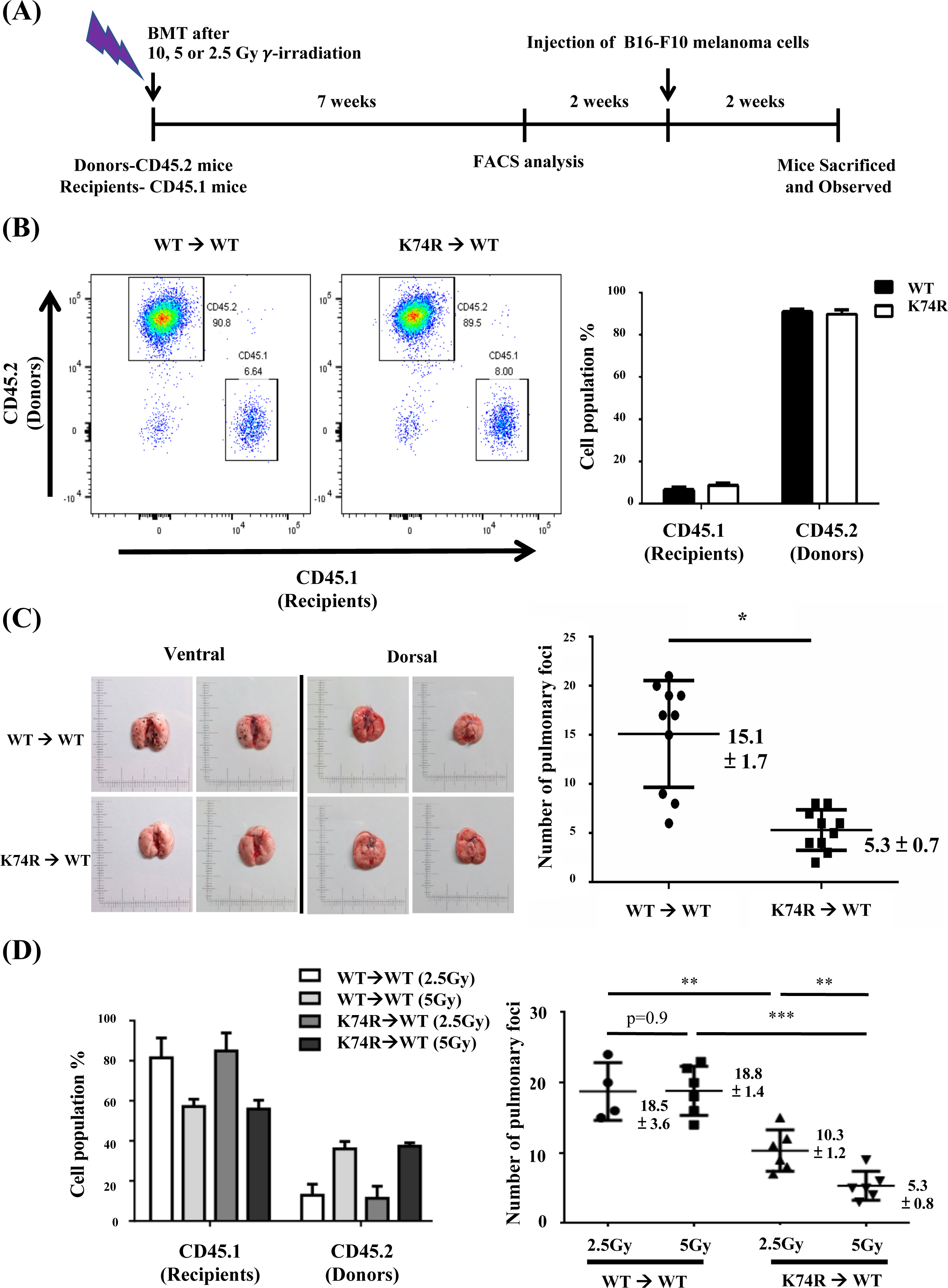
Transfer of cancer resistance of *Klf1*(K74R) mice to WT mice by bone marrow transplantation (BMT) (A) Flow chart illustrating the experimental strategy. (B) FACS analysis of the efficiency of BMT with use of 10Gy γ-irradiation. The percentages of CD45.1/CD45.2 cells in the PB of the recipient male mice were analyzed by flow cytometry, with the representative FACS charts shown on the left and the statistical histobar diagram on the right. (C) Transfer of the anti-metastasis capability of 8 week-old *Klf1*(K74R) male mice to age-equivalent WT male mice by BMT with use of 10Gy γ-irradiation. Left panels, representative photographs of lungs with pulmonary metastatic foci in the recipient WT (CD45.1) mice after BMT from WT (CD45.2) or *Klf1*(K74R) (CD45.2) donor mice and challenged with B16-F10 cells. Statistical analysis of the numbers of pulmonary B16-F10 metastatic foci on the lungs is shown in the right histogram. n=10, *, p<0.05. (D) Transplantation of 8 week-old male WT (CD45.1) mice with BMMNC from age-equivalent WT (CD45.2) male mice or from *Klf1*(K74R) (CD45.2) male mice with use of the γ-irradiation dosage 2.5Gy or 5Gy. The histobar diagram comparing the percentages of CD45.1 and CD45.2 PB cells of the recipient WT mice after BMT is shown on the left. The statistical analysis of the average numbers of pulmonary foci on the lungs of recipient WT mice after BMT and injected with the B16-F10 cells is shown in the right histogram, N=6. **, p<0.01, ***, p<0.001.

In order to determine if WT mice having more restricted blood replacement upon BMT from *Klf1*(K74R) mice still exhibited a higher anti-cancer capability, we also carried out BMT experiments with lower doses of γ-irradiation. BMT using two lower doses of γ-irradiation (2.5Gy/5Gy) still resulted in transfer of cancer resistance from *Klf1*(K74R) to WT mice. Approximately 40% of recipient blood cells were substituted by donor cells upon BMT with 5Gy γ-irradiation. Consequently, at that irradiation dosage, BMT from *Klf1*(K74R) mouse donors reduced the average number of pulmonary tumor foci in recipient WT mice to 5. On the other hand, only 20% blood replacement was achieved in the recipient mice with 2.5Gy γ-irradiation (Figure 2D). However, the WT mice receiving BMT from *Klf1*(K74R) mice again developed less number (∼10/mouse) of pulmonary tumor foci than those WT mice receiving BMT from the WT mice (Figure 2D). Thus, even at a low level of 20% blood replacement, BMT still enabled effective transfer of cancer resistance from *Klf1*(K74R) mice to WT mice.

In addition, we also attempted to transfer the extended lifespan characteristics of the *Klf1*(K74R) mice to WT mice by BMT. Significantly, the median lifespan of WT mice receiving BMT from *Klf1*(K74R) mice was 5 months longer than that of WT mice receiving BMT from WT mice (Figure 2-figure supplement 1B). Thus, the longer lifespan characteristics of the *Klf1*(K74R) mice was also transferable via BMT.

### Inhibition of tumor growth by transplanted BMMNC from *Klf1* K74R) mice

Our experiments indicated that *Klf1*(K74R) bone marrow carried the anti-metastasis capability that prevented melanoma cell colonization on the lungs of recipient mice (Figures. 1 and 2). To determine if *Klf1*(K74R) BMT could inhibit tumor growth, we examined the effect of BMT on growth of tumors with B16-F10-luc cells. As outlined in Figure 3A, ten days after injection of cancerous cells, the formation of bioluminescent signals in the recipient mice were confirmed by the observation of *in vivo* bioluminescence. The following day, we transplanted the recipient mice with BMMNC from WT or *Klf1*(K74R) mice and then measured the intensities of bioluminescence signals from tumors 7 and 14 days later. As shown, tumor growth in mice subjected to BMT from *Klf1*(K74R) mice was significantly slower relative to those receiving BMMNC from WT mice (Figure 3B and 3C). Thus, *Klf1*(K74R) BMMNC indeed can inhibit the growth of tumor more effectively than WT BMMNC.

**Figure 3.**
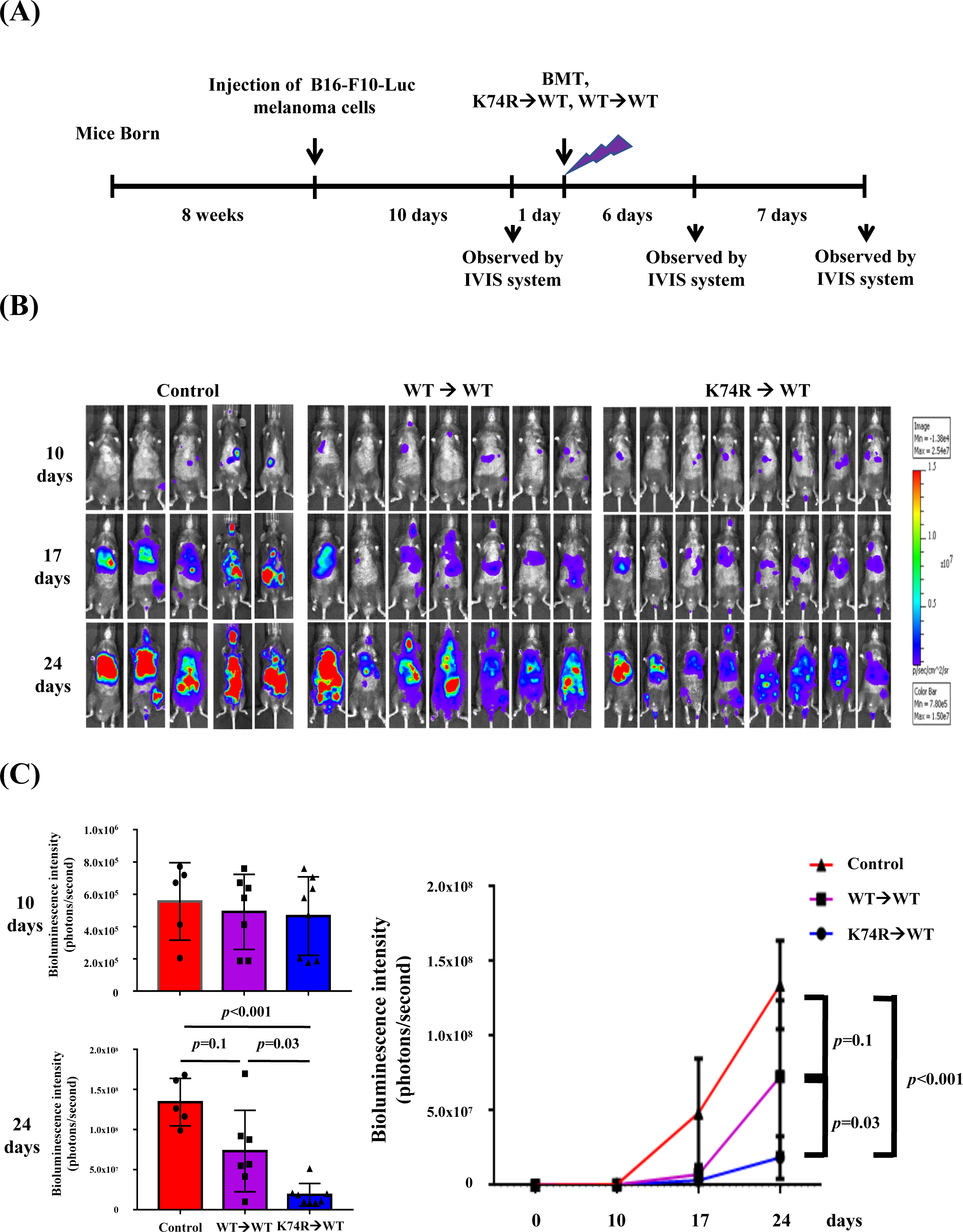
Inhibition of tumor growth in WT mice by BMT from *Klf1*(K74R) mice. (A) A flow chart of the experiments. Luciferase-positive B16-F10 cells were injected into the tail vein of 8 week-old WT male mice (day 0). The mice were then transplanted with BMMNC from WT or *Klf1*(K74R) male mice on day 11 after the luciferase-positive B16-F10 cell injection. *In vivo* imaging system (IVIS) was used to follow the tumor growth in mice on day 0, 10, 17 and 24, respectively. (B) Representative images of bioluminescence reflecting the luciferase activity from melanoma cancer cells in mice. The color bar indicates the scale of the bioluminescence intensity. (C) Statistical analysis of the intensities of bioluminescence in the cancer-bearing mice (WTèWT, purple, N=7; *Klf1*(K74R)èWT, blue, N=8; Control (no BMT), red, N=3).

### Differential expression of specific immune-, ageing- and/or cancer- associated biomolecules in the blood of *Klf1*(K74R) mice

In general, the cell populations in PB were not affected much by the K74R substitution, as shown by CBC analysis^21^, although flow cytometry analysis of PB from WT and *Klf1*(K74R) mice of different ages showed that the population of NK(K74R) cells in 24 month-old *Klf1*(K74R) mice was higher than WT NK cells in 24 month-old WT mice (Figure 4-figure supplement 1). In parallel, the K74R substitution did not affect much the expression of *Klf1* gene in the different types of hematopoietic cells. As seen in Figure 4-figure supplement 2, the K74R substitution did not alter the expression levels of KLF1 protein in leukocytes from peripheral blood (PB) (left panels, Figure 4-figure supplement 2A), although the *Klf1* mRNA level of *Klf1*(K74R) leukocytes was 10-20% lower than that of WT leukocytes (right panel, Figure 4-figure supplement 2A). Previous studies and database have shown that *Klf1* mRNA and KLF1 protein are expressed in the erythroid lineage^10^ and megakaryocytes^10,14^, and the *Klf1* mRNA is expressed in HSC^16^(bio-GPS database), NK cells (bio-GPS database), naïve T cells^12^ and Treg cells^12^. By Western Blotting (WB) analysis (Figure 4-figure supplement 2B) and RT-qPCR analysis (Figure 4-figure supplement 2C), we found that the *Klf1* gene is expressed in B220^+^ B cells, CD3^+^ T cells, and NK cells of both WT and *Klf1*(K74R) mice, but the levels are much lower than that of the erythroid MEL cells.

**Figure 4.**
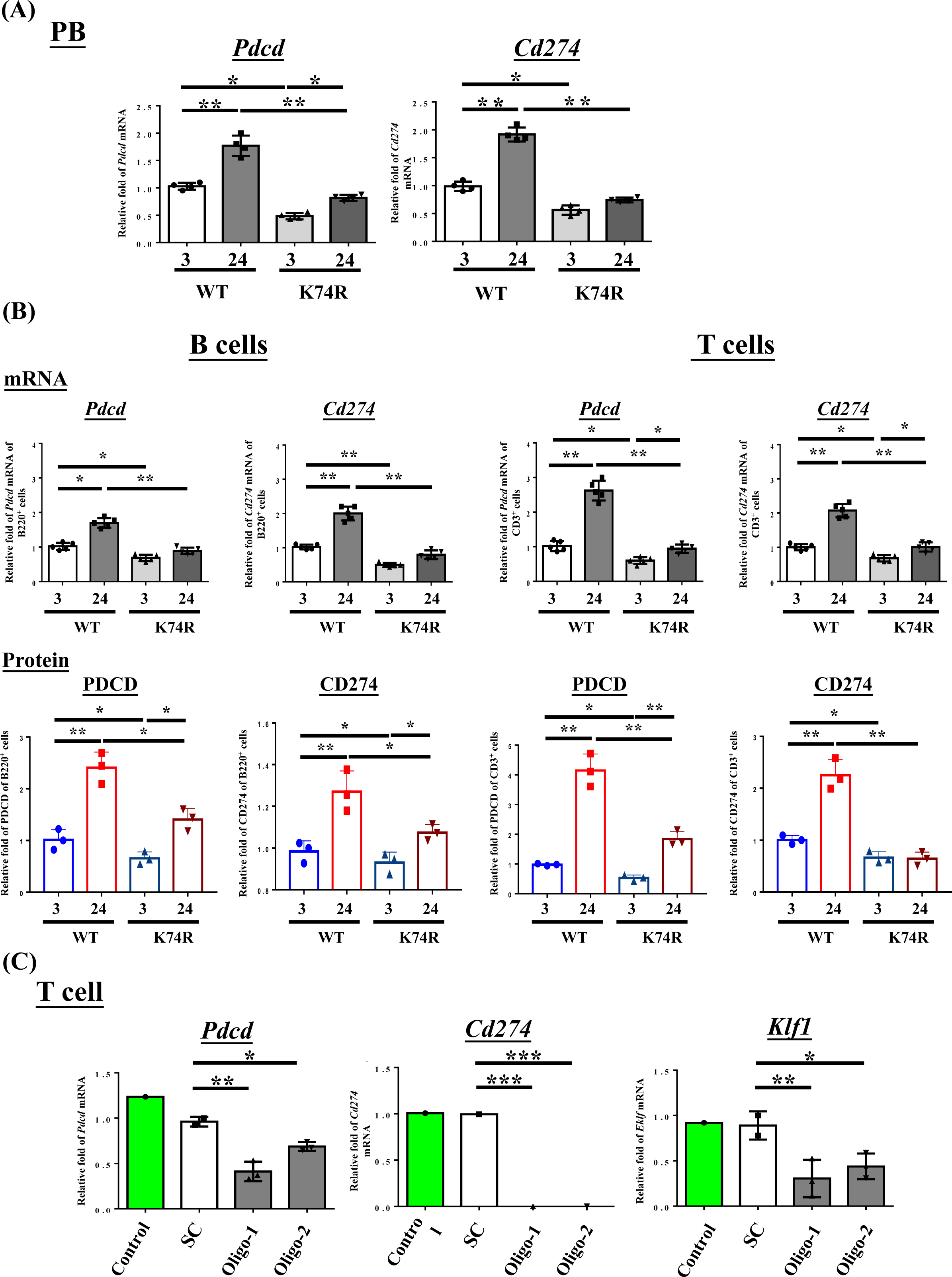
Decrease of *Pdcd* and *Cd274* expression in blood cells of *Klf1*(K74R) mice. (A) Levels of *Pdcd* and *Cd274* mRNAs in the PB of WT and *Klf1*(K74R) male mice at the ages of 3 months and 24 months, respectively, as analyzed by RT-qPCR. Note the relatively low levels of *Pdcd* and *Cd274* mRNAs in the *Klf1*(K74R) mice at both ages in comparison to the WT mice. (B) Upper panels, comparison of the mRNA levels of *Pdcd* and *Cd274* of CD3^+^ T cells and B220^+^ B cells isolated from the PB of 8 week-old WT and *Klf1*(K74R) male mice. N=5. *, p<0.05; **, p<0.01. Lower panels, comparison of the protein levels of PDCD and CD274, as analyzed by flow cytometry, of CD3^+^ T cells and B220^+^ B cells from 8 week-old WT and *Klf1*(K74R) male mice. N=3. *, p<0.05; **, p<0.01. (C) Comparison of the levels of *Pdcd, Cd274* and *Klf1* mRNAs, as analyzed by RT-qPCR, in CD3^+^ T cells, which were isolated from splenocytes, without or with RNAi knockdown of *Klf1* mRNA. Two oligos (oligo-1 and oligo-2) were used to knockdown *Klf1* mRNA by ∼60-70%, which resulted in the reduction of *Pdcd* mRNA level by 30-60% and nearly complete depletion of *Cd274* mRNA. Control, T cells transfected with GFP-plasmid. SC, T cells transfected with scrambled oligos. N>3. *, p<0.05; **, p<0.01; ***, p<0.001.

We first used RT-qPCR to analyze the levels in PB cells of mRNAs encoding the immune checkpoint genes (ICGs) PDCD/CD274^25^ in view of the cancer resistance of *Klf1*(K74R) mice (Figure 4) as well as increased levels of PDCD and CD274 in aged or tumorigenic mice^26^. As shown in Figure 4, the mRNA levels of *Pdcd* and *Cd274* in the PB, B cells and T cells of WT mice were both increased during aging. In great contrast, the mRNA levels and protein levels of these two genes were lower in 3 month-old *Klf1*(K74R) mice than the age-matched WT mice, and they remained low during ageing of the *Klf1*(K74R) mice. Significantly, RNAi knockdown of *Klf1* expression reduced *Pdcd* and *Cd274* mRNA levels in splenic CD3^+^ T cells (Figure 4C). In interesting correlation, overexpression of KLF1 would increase the expression of *Cd274* in mouse CD4^+^ T cells^12^. There data together indicate that KLF1 positively regulates the transcription of the *Pdcd* and *Cd274* genes, directly or indirectly, and likely the transcriptomes of a diverse range of hematopoietic cells. Since the levels of KLF1 protein in CD3^+^ T (K74R) cells and WT CD3^+^ T cells were similar (Figure 4-figure supplement 2B), change of the expression of *Pdcd* and *Cd274* genes (Figure 4) was most likely caused by the loss of the sumoylation of KLF1(K74R), which would alter its transactivation capability^17^. Interestingly, *Pdcd* appeared to be a direct regulatory target of KLF1. As shown by ChIP-qPCR analysis, KLF1 was bound at the CACCC box (Box3) at −103 of the *Pdcd* promoter in CD3^+^ T cells from both WT and *Klf1*(K74R) mice (Figure 4-figure supplement 3). The detail of the role of sumoylation in the transcriptional activation of *Pdcd* by KLF1 awaits future investigation. As expected, lower levels of *Pdcd* and *Pdcd1* expression were also observed in the PB of mice receiving BMT from *Klf1*(K74R) mice (Figure 4-figure supplement 2D). These findings together suggest that the low tumorigenesis rate of *Klf1*(K74R) mice arises in part from low expression of the two ICGs, *Pdcd* and *Cd274*, as the result of K74R substitution in the KLF1 protein.

We have also examined, by bead-based multiplex assay^27,28^, the expression patterns of several ageing- and/or cancer-associated cytokines. As shown in Figure 4-figure supplement 4, there was no significant difference in the serum levels of IL-1β, IL-2, IL-10, IL-12p70, INF-γ or TNF-α between WT and *Klf1*(K74R) mice at 24 month-old. In contrast, the level of IL-4, an anti-inflammatory cytokine^29^ beneficial to the hippocampus of aging mice^30^, in 24 month- old *Klf1*(K74R) mice was 3-4 fold higher than the 24 month-old WT mice. On the other hand, the level of IL-6, a key factor in chronic inflammatory diseases, autoimmunity, cytokine storm and cancer^28,31^, increased only moderately during aging of the *Klf1*(K74R) mice (Figure 4-figure supplement 4). Thus, similar to PDCD and CD274, the altered expression of some of the cytokines in the blood likely also contributes to the anti-aging and/or anti-cancer characteristics of the *Klf1*(K74R) mice.

### Comparative proteomics analysis of the leukocytes of *Klf1*(K74R) mice and WT mice

We proceeded to examine age-dependent cell-intrinsic changes in the proteomes of the leukocytes from the WT and *Klf1*(K74R) mice in two different age groups. 259 and 306 differentially expressed proteins (DEPs) were identified between the two age groups for the WT and *Klf1*(K74R) mice, respectively (Figure 5A). To understand the correlations of these proteins with aging and cancer, we performed the GSEA and found that the age-dependent DEPs changed in the concordant direction in the WT and *Klf1*(K74R) mice were enriched for several known aging-related pathways, e.g. IL-6-JAK-STAT3 signaling, DNA repair, etc^32^ (Figure 5B). Meanwhile, the age-dependent DEPs changed in the reverse directions in the WT and *Klf1*(K74R) mice were enriched for nine other aging-related pathways (Figure 5C).

**Figure 5.**
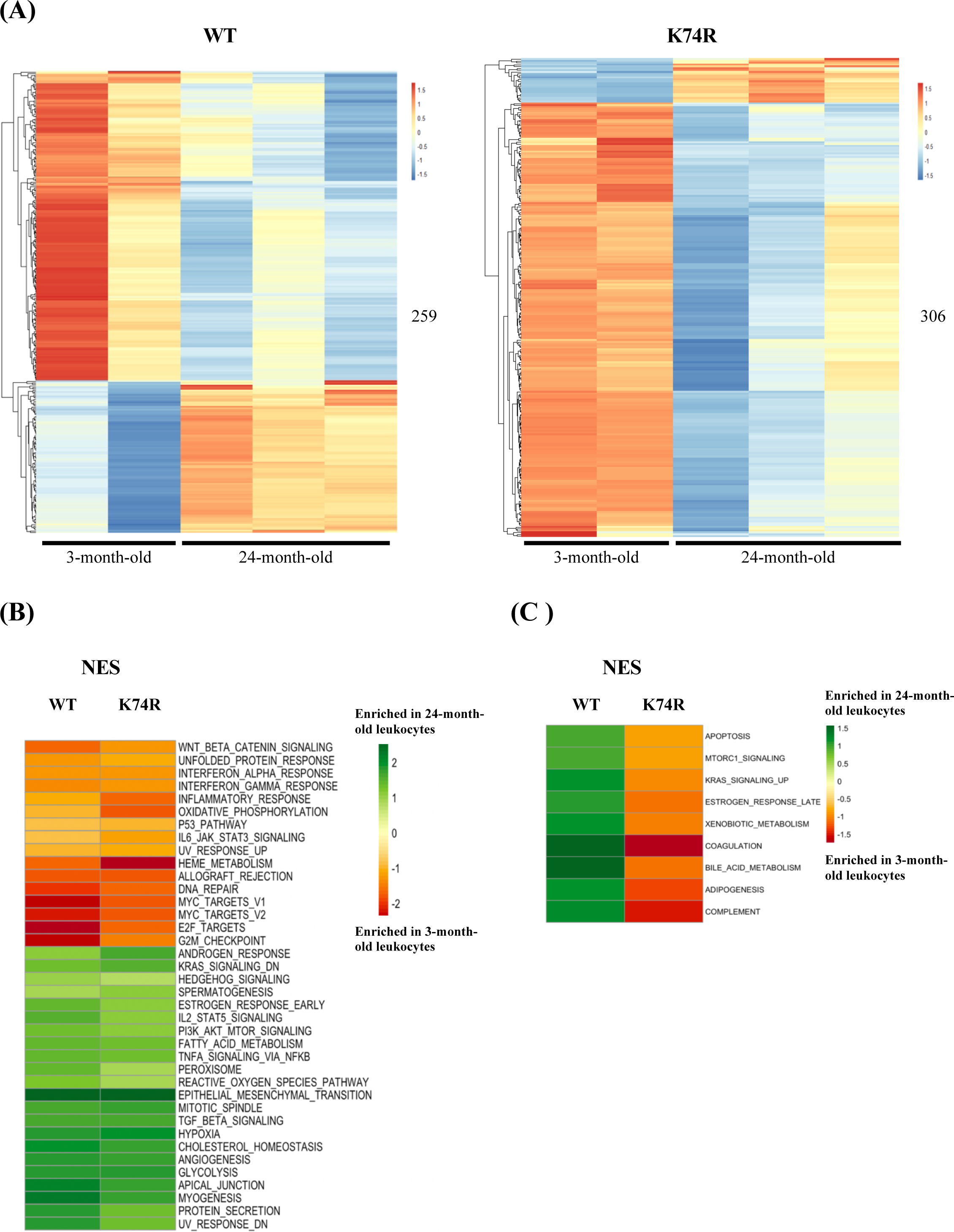
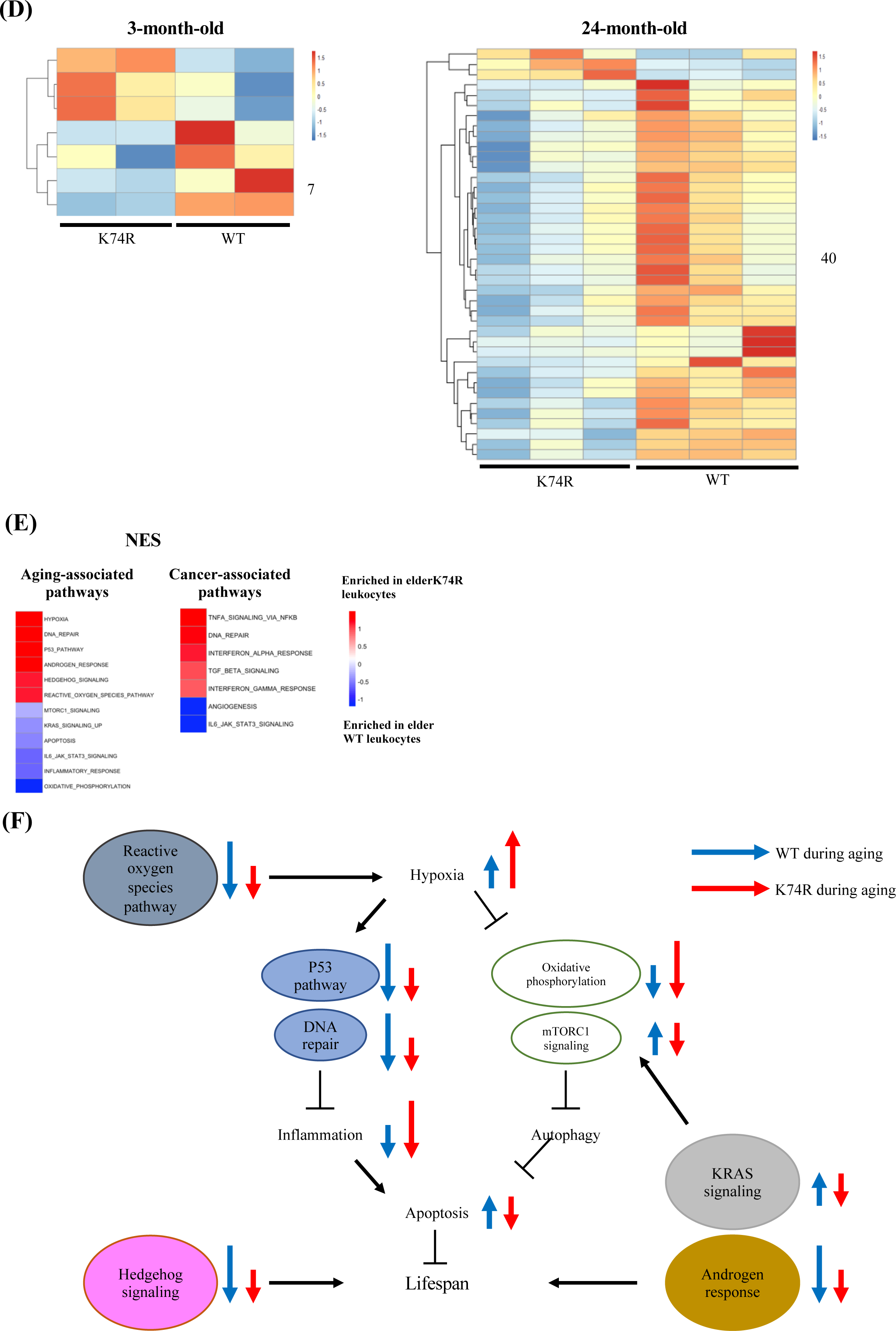
Proteomics analysis of mouse leukocytes. (A) Heatmap plots representing age-dependent differentially expressed proteins (DEPs) in the leukocytes from the WT male mice (left) and *Klf1*(K74R) male mice (right). Differential Test Threshold: expression fold change > 1.5 and *p*-value < 0.01. (B) and (C), pathway analysis of the age-dependent DEPs changed in the concordant (B) and reverse (C) directions, respectively, in the WT and *Klf1*(K74R) mice. NES, normalized enrichment score. (D) Heatmap plots representing strain-dependent DEPs in leukocytes of 3 month-old (left) and 24 month-old (right) mice. (E) Pathway analyses of the strain-dependent DEPs in elder WT leukocytes (enrichment indicated by blue color) and elder *Klf1*(K74R) leukocytes (enrichment indicated by red color). The aging- and cancer-associated pathways are presented in the left and right diagrams, respectively. NES, normalized enrichment score. (F) A model of the regulation of the mouse lifespan by different cellular pathways in the leukocytes. The inter-relationship of 11 aging-associated cellular pathways differentially expressed in the leukocytes of *Klf1*(K74R) male mice in comparison to the WT male mice, among themselves and with respect to the regulation of lifespan (for references, see text), are depicted here. The directions and lengths of the arrows indicate changes of the individual pathways, up-regulation (upward arrows) or repression (downward arrows), in the leukocytes during aging of the WT male mice (blue arrows) and *Klf1*(K74R) male mice (red arrows), respectively.

We further performed DEP analyses between WT and *Klf1*(K74R) mice and identified strain-dependent DEPs in the two age groups. As shown in Figure 5D, only 7 DEPs were identified in the 3 month-old mice but 40 DEPs in the 24 month-old ones. Of the 40 DEPs in the elder mice, 3 and 37 were upregulated in *Klf1*(K74R) and WT mice, respectively (Figure 5D). Significantly, GSEA analysis of these DEPs showed that elder *Klf1*(K74R) leukocytes were enriched for the anti-aging pathways related to hypoxia, and p53 signaling, etc.^33,34^, while the elder WT leukocytes were enriched for the aging-associated pathways related to apoptosis, and mTORC1 signaling, etc^33,35^. On the other hand, the DEPs in elder *Klf1*(K74R) leukocytes were also enriched for anti-cancer pathways related to the interferon-α response, and TGF-β signaling, etc.^36–38^ (Figure 5E), while DEPs in the elder WT leukocytes were enriched for the pro-cancer pathways related to IL-6-JAK-STAT3 signaling and angiogenesis^39^. These data together have demonstrated that *Klf1*(K74R) leukocytes contribute to their anti-cancer capability and long lifespan through several specific cellular signaling pathways.

### Higher *in vitro* cancer cell cytotoxicity of the NK(K74R) cells than WT NK cells

The higher anti-cancer capability of the *Klf1*(K74R) mice could result from combined contributions by different types of the hematopoietic cells, e.g. the NK(K74R) cells. NK cells are effector lymphocytes of the innate immune system that can rapidly destroy tumor cells and against infectious pathogen^40^. Also, NK cells do not require pre-stimulation for the treatment of cancer cells^41^. Further, the NK lineage could be fully reconstituted 2 weeks after BMT^42^. We tested the *in vitro* cancer cell cytotoxicity ability of NK(K74R) cells from 3 month-old *Klf1*(K74R) mice in comparison to the WT NK cells. As shown in Figure 6, NK(K74R) cells exhibited significantly higher cytotoxicity against the B16-F10 melanoma cells as well as Hepa1-6 hepatocellular carcinoma cells than WT NK cells. This data together with the higher level of NK cells in aged *Klf1*(K74R) mice (Figure 4-figure supplement 1) indicated that in parallel to the leukocytes^21^, NK cells also play an important role in the higher anti-cancer capability of the hematopoietic blood system of *Klf1*(K74R) mice, as exemplified in Figures 2 and 3.

**Figure 6.**
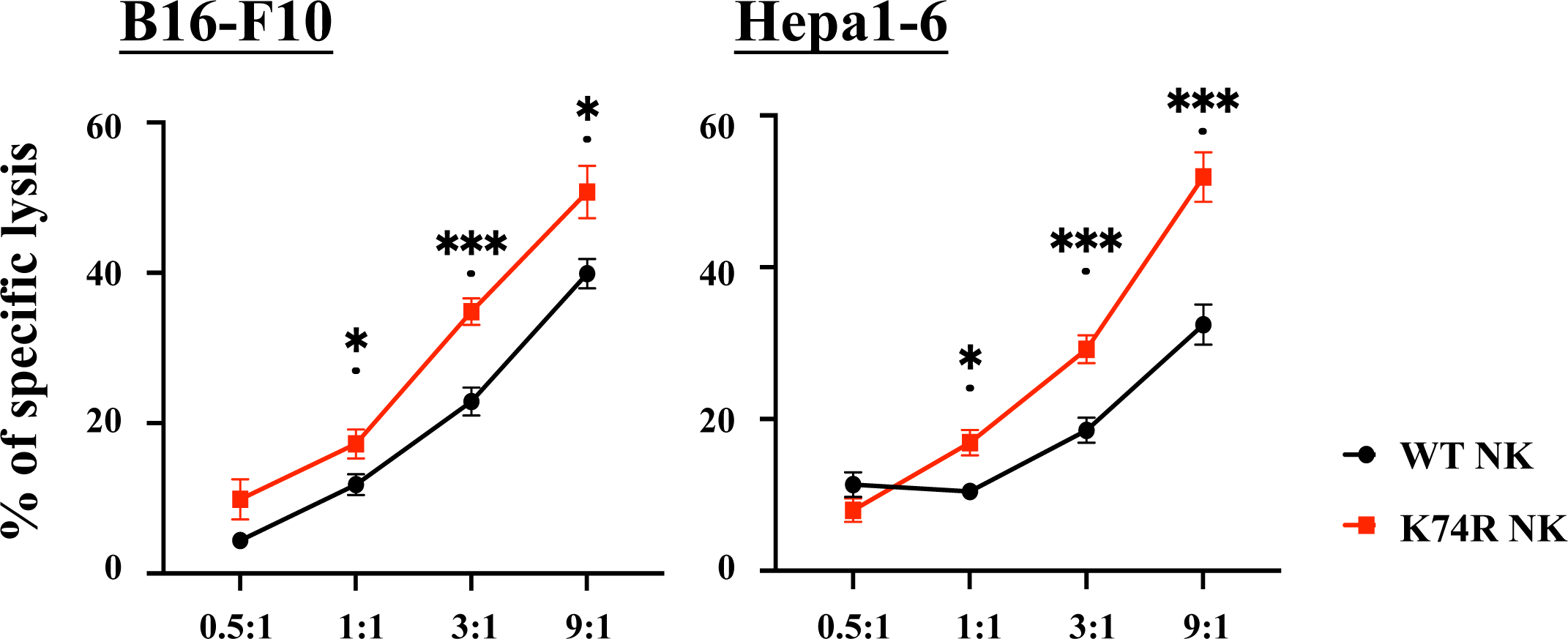
Higher *in vitro* cancer cell cytotoxicity of NK(K74R) cells than WT NK cells. B16-F10 and Hepa1-6 cells were labeled with Calcein-AM (BioLegend) and co-cultured with NK cells in different E:T ratio for 4hrs. Killing rates of B16-F10 and Hepa 1-6 cancer cells by NK cells were determined by the intensities of fluorescence in the medium in comparison to the controls of spontaneous release and maximwal release.

## Discussion

Because of the complexity and intercrosses of the different pathways regulating the health and the aging process, genetic manipulation of non-human animals^43,44^ and non-genetic approaches on animals including human^4,45,46^ targeting these pathways inevitably lead to moderate-to-severe side effects such as body weight loss, adiposity, etc.. With respect to the above, the *Klf1*(K74R) mice^21^ is ideal as an animal model for further insightful understanding of the ageing process as well as for biomedical development of new anti-ageing tools and approaches. In the current study, we demonstrate that the healthy longevity of the *Klf1*(K74R) mice, as exemplified by their higher tumor resistance, is likely to be gender/genetic background independent and irrespective of the amino acid change at K74 of KLF1. Further, we show that the *Klf1*(K74R) mice are resistant to carcinogenesis of not only melanoma, but also other cancers such as hepatocellular carcinoma. This correlates well with the higher cytotoxicity *in vitro* of NK(K74R) cells than WT NK cells against the same two types of cancer cells. Significantly, we also show that one-time transplantation of the bone marrow mononuclear cells or BMMNC (BMT) could confer the recipient WT mice extended lifespan and higher tumor-resistance capability, the latter of which could be achieved by 20% of substitution of the blood of the recipient WT mice with that from the *Klf1*(K74R) mice. These findings together have interesting implications in the development of this novel hematopoietic blood system for future medical application in anti-disease and anti-aging.

The anti-cancer capability of the *Klf1*(K74R) mice have rendered them relatively free from spontaneous cancer occurrence^21^, which is also reflected by their resistance to tumorigenesis of the B16-F10 cells and Hepa1-6 in the anti-cancer assays (Figures 1-3). Furthermore, the cancer resistance of *Klf1*(K74R) mice appears to be independent of the gender, age, and genetic background (Figure 1). The anti-metastasis property of the *Klf1*(K74A) mice in the pulmonary foci assay (Figure 1D) also indicates that the anti-cancer capability of the *Klf1*(K74R) mice is not due to the structural and/or post-translational properties of the arginine introduced at codon 74 of KLF1. Importantly, we have shown that the anti-cancer capability and the extended lifespan characteristics of *Klf1*(K74R) mice are transferrable through BMT (Figure 2, 3, and S2). In particular, we show that BMMNC from *Klf1*(K74R) mice (Figure 2 and S2A) could confer 2 month-old WT recipient mice with the anti-cancer capability against melanoma. Furthermore, ∼20% of blood substitution would allow the recipient mice to become cancer-resistant in the pulmonary foci assay (Figure 2D). It is likely that homozygosity, but not heterozygosity, of K74R substitution causes one or more types of the hematopoietic cells to gain new functions, such as the higher cancer cell-killing capability of NK(K74R) cells (Figure 6). The data of Figure 2D further suggest that the amount of specific types of cells such as NK(K74R) in the 20% of blood cells carrying homozygous *Klf1*(K74R) alleles in the recipient mice upon BMT is sufficient to confer the mice a higher anti-cancer ability.

Hematopoietic stem cell therapy for different diseases^47–50^ including cancer has been intensively explored and practiced such as leukemia, and neuroblastoma, etc^51,52^. Also, certain characteristics of the young mice could be transferred to old mice via heterochronic parabiosis or heterochronic transplantation^53–56^. Similarly, plasma proteins from human umbilical cord blood can revitalize hippocampal function and neuroplasticity in aged mice^57,58^. Previously, we have shown that multiple injections of LSK-HSC(K74R) from 3 month-old *Klf1*(K74R) mice to 25 month-old WT mice would extend the lifespan of the latter by about 4 months^21^. In the current study, we further show that 2 month-old WT mice receiving one-time transplantation of BMMNCs from 2 month-old *Klf1*(K74R) mice would live 5.5 months longer than the controls (Figure 2-figure supplement 1B). Thus, transplantation/transfer of the blood MNC carrying homozygous mutation at the sumoylation site of KLF1 could be developed as a new approach for long-term anti-aging, anti-cancer and likely rejuvenation.

The tumorigenesis resistance and long lifespan exhibited by the *Klf1*(K74R) mice are most likely due to changes in the transcription regulatory properties of the mutant KLF1(K74R) protein relative to WT^59^. As exemplified in Figure 4A and 4B, expression levels of the ICGs *Pdcd* and *Cd274* in the PB, B, and T cells of *Klf1*(K74R) mice are reduced in comparison to the WT mice. Notably, cancer incidence increases with aging^60^, which is accompanied by increased expression of PDCD and CD274^61^. The lower expression of ICGs would contribute to the anti-cancer capabilities of the *Klf1*(K74R) blood to fight against cancer (Figures. 1, 2, and 3) and to extension of the lifespan of cancer-bearing mice^21^. Indeed, similar to ICGs, the expression levels of several cytokines in the *Klf1*(K74R) blood/serum are also different from the WT blood and some of the changes during ageing or carcinogenesis in the *Klf1*(K74R) blood are opposite to the blood/serum of WT mice ^28^ (Figure 4-figure supplement 4).

The transcriptome data have been used to dissect the regulation of leukocyte aging ^32,62,63^. In addition, proteomic analysis has revealed the signaling pathways that regulate aging of specific types of leukocyte such as the lymphocyte and neutrophils cells^64,65^. In this study, we have performed proteomics analysis of leukocytes from WT and *Klf1*(K74R) mice in two age groups, and found that for the elder mice, the strain-dependent DEPs in the leukocytes are enriched for a number of signaling pathways. Among these signaling pathways, at least 12 of them are closely associated with the aging process, which include hypoxia, DNA repair, etc. (Figure 5E). As summarized by the model in Figure 5F, it appears that changes of these pathways in the elder *Klf1*(K74R) leukocytes relative to the elder WT leukocytes are mostly in the direction of anti-aging. The data of Figure 5 together strongly suggest that the KLF1(K74R) amino acid substitution causes a change in the global gene expression profile of the leukocytes, which contributes to the high anti-cancer capability and long lifespan of the *Klf1*(K74R) mice. In sum, we have characterized the cancer resistance of the *Klf1*(K74R) mice, among their other healthy characteristics, in relation to gender, age, and genetic background. We also have identified cell populations, gene expression profiles and cellular signaling pathways of the white blood cells of young and old mutant mice, in comparison to the WT ones, that are changed in the anti-cancer and/or anti-ageing directions. Finally, the transferability of the cancer resistance and extended life-span of the mutant mice via transplantation of BMMNC suggests the possibility of future development of hematopoietic blood cells genome-edited at the conserved sumoylation site of KLF1 for anti-cancer and the extension of healthspan and/or lifespan in animals including human.

## Acknowledgments

We thank Ya-Min Lin and her colleagues at the FACS Core for their efforts on the cell sorting. We also thank Taiwan Mouse Clinic facility for expertise help on the IVIS experiments. The effort by Dr. Ching-Yen Tsai at the Transgenic Core Facility, IMB, is greatly appreciated. The Immune Monitoring Core of Taipei Medical University provided great help in the analysis of the blood cell populations. We also thank the Bioinformatics Core Facility, IMB, for its efforts on the proteomics analysis, and Dr. Da-Liang Ou at National Taiwan University for providing us the Hepa1-6 cell line.

## Animal

All the animal procedures were approved by the Institute of Animal Care and Use Committees, IACUC, at Academia Sinica. The IACUC numbers are 17-02-1052, 17-12-1142, 20-10-1528, BioTReC-110-M-005, and BioTReC-111-M-004, respectively.

## Availability of data and materials

The data that support the findings of this study are available from the corresponding authors upon request.

## Material and Methods

### Mice

C57BL/6, B6, and FVB mice were purchased from Jackson Laboratories (Bar Harbor, Maine). The B6 *Klf1*(K74R), B6 *Klf1*(K74A) and FVB *Klf1*(K74R) mice were established with the assistance of the Transgenic Core Facility (TCF), IMB, Academia Sinica, Taiwan. As described previously^21^, the K74R mutation was introduced by homologous recombination into exon 2 (E2) of the *Klf1* gene of B6 mice by means of a recombinant retrovirus containing the construct loxP-PGK-gb2-neo-loxP-E2 (K74R), before excising the neomycin (neo) selection marker by crossing with Ella-Cre mice. The heterozygous *Klf1*(K74R/+) mice were then crossed to obtain homozygous mutant *Klf1*(K74R/K74R) mice, hereafter termed *Klf1*(K74R) mice.

On the other hand, *Klf1*(K74A) mice were generated by using the CRISPR/Cas9 system. Female B6 mice (7 to 8 week-old) were mated with B6 males and the fertilized embryos were collected from the oviducts. For oligos injection, Cas mRNA (100 ng/µl), sgRNA (50 ng/µl), and donor oligos (100 ng/µl) were mixed and injected into the zygotes at the pronuclei stage. The F0 mice were genotyped by PCR and DNA sequencing. The heterozygous *Klf1*(K74A/+) mice were crossed to establish the germ-line stable homozygous *Klf1*(K74A) F1 strain.

*Klf1*(K74R) mice in the FVB background were generated using an *in vitro* fertilization strategy. Briefly, sperm from male B6 *Klf1*(K74R) mice was used to fertilize FVB mouse oocytes. *In vitro* fertilizations of FVB oocytes were carried out consecutively for five generations. The resulting chimeric mice with >90% FVB background were then crossed with FVB mice for another five generations or more.

### Cell lines

Murine B16-F10 melanoma cell line was purchased from ATCC (CRL-6475). B16-F10 cells expressing luciferase (B16-F10-luc) were generated by infection of recombinant lentivirus stably expressing luciferase luc2 under the control of hEF1A-eIF4G promoter (InvivoGen)^61^. All cell lines were derived from cryopreserved stocks split fewer than three times and they were confirmed as mycoplasma-free prior to use. B16-F10 cells were cultured at 37 °C and 5 % CO_2_ in DMEM medium supplemented with 10 % FBS, 1 % penicillin/streptomycin, and 2 mM L-glutamine. B16-F10-luc cells were selected at 37 °C and 5 % CO2 in a DMEM medium supplemented with 0.2 mg/mL zeocin (Invitrogen), 10 % FBS, 1 % penicillin/streptomycin, and 2 mM L-glutamine. The hepatocellular carcinoma cell line, Hepa1-6, was from Dr. Da-Liang Ou’s lab at College of Medicine, National Taiwan University. The line was maintained at 37°C and 5% CO_2_ in Gibico^TM^ DMEM, a high glucose commercial medium containing 10% FBS (Gibico^TM^) and 1% penicillin-streptomycin.

### Pulmonary melanoma foci assay

Cultured B16-F10 melanoma cells (1×10^5^ cells/mouse) were injected into the tail vein of 8 week- to 9 week-old or 24 month-old *Klf1*(K74R) and WT mice with/ without bone marrow transplantation. Two weeks after injection, the mice were sacrificed and the number of tumor foci on their lungs was quantified^66^.

### Subcutaneous cancer cell inoculation assay of Hepa1-6

The protocol follows that used by Han *et al*^67^. After anaesthetization of the mice with 3% isoflurane, 2×10^6^/ 50uL of Hepa1-6 cells in serum-free DMEM medium were mixed with 50ul of Corning® Matrigel® Matrix and injected between skin and muscle of the hind limb. The size of the tumors was measured and recorded once a week with Mitutoyo^®^ Vernier Caliper. The volume was calculated using the formula: length × weight × height × π/6.

### Flow cytometric analysis and cell sorting

Single cell suspensions of the peripheral blood cells and spleen tissue of B6 mice were prepared by lysing red blood cells and then passing them through a 40-µm cell strainer (Falcon®). Bone marrow mononuclear cells (BMMNCs) were prepared as described below in the section **Bone marrow transplantation (BMT)**. The peripheral blood cells and splenocytes were stained extracellularly for 30 min at room temperature using different combinations of the following antibodies: anti-CD45.1 (eBioscience); anti-CD45.2 (eBioscience); anti-CD3e (eBioscience); anti-CD45R (eBioscience); anti-NK1.1 (eBioscience); anti-NKp46 (eBioscience); anti-Gr-1 (eBioscience); CD11b (eBioscience); anti-PDCD (eBioscience) and anti-CD274 (eBioscience). The various hematopoietic progenitor cell compartments of bone marrow were also stained extracellularly for 30 min at room temperature by using different combination of the following antibodies: anti-Lineage (eBioscience), anti-c-Kit (eBioscience), anti-Sca-1 (eBioscience), anti-CD34 (eBioscience), anti-Flt-3 (eBioscience). All the immuno-stained cells were subsequently washed with 1% PBS three times and resuspended for FACS analysis and sorting. Small amounts of the cell samples were run on a FACS Analyzer LSRII-12P (BD Bioscience) to determine the proportions of different cell preparations. FACS AriaII SORP (BD Bioscience) was then used to sort the indicated cell populations. The detail gating subsets for all the cell as described above are shown in Supplementary file 1a. Data analysis was performed using FlowJo software.

### RNAi knockdown of *Klf1* mRNA from T cells

CD3^+^ T cells isolated by sorting were cultured in RPMI 1640 medium for one day for recovery and then transfected with EGFP-plasmid (control), scrambled oligonucleotides (SC control), *Klf1* knockdown oligonucleotide 1 (oligo 1), or *Klf1* knockdown oligonucleotide 2 (oligo 2) in a 96-well plate for 48 h using a LONZA electroporation kit (P3 Primary Cell 4D-NucleofectorTM X Kit) and machine (4D-NucleofectorTM Core Unit). Then the cells were lysed using a PureLink® RNA Mini kit (Life Technologies) and analyzed by RT-qPCR.

### RT-qPCR analysis

Total RNAs of B cells, T cells, and leukocytes/ WBC from the peripheral blood or spleen of *Klf1*(K74R) mice and WT mice, respectively, were extracted using a PureLink® RNA Mini kit (Life Technologies). The RNAs were reverse-transcribed by means of oligo-dT primers, Maxima H Minus Reverse Transcriptase (Thermo Scientific™) and SYBR Green reagents (Applied Biosystems). RT-qPCR was performed using a LightCycler® Nano machine (Roche). Gene-specific primers for *Klf1*, *Pdcd*, *Cd274* and *Gapdh* were designed using Vector NTI Advance 9 software according to respective mRNA sequences in the NCBI database (primer sequences are available upon request). The PCR primers for analysis of *Klf1* mRNA had been used and verified in a previous study^16^. Expression levels of mRNAs were normalized to that of endogenous *Gapdh* mRNA.

### ChIP-qPCR

The ChIP-qPCR analysis followed the procedures by Daftari *et al*^68^. Extracts were prepared from formaldehyde cross-linked mouse CD3^+^ T cells and erythroleukemia cell line (MEL), sonicated, and then immune-precipitated (IP) with anti-KLF1 (Thermo) or purified rabbit IgG. The chromatin DNAs in the precipitates were purified and analyzed by qPCR in the Roche LightCycle real-time system. Sequences of the DNA primers used for qPCR are shown in Supplementary file 1b.

### Western blotting (WB)

We adopted a previously described WB procedure^69^. Leukocytes/ WBC of *Klf1*(K74R) mice and WT B6 mice, respectively, were collected from RBC lysis buffer-treated peripheral blood. CD3^+^ T cells, B220^+^ B cells and NK cells from the spleen were sorted by FACS AriaII SORP (BD Bioscience). The cell pellets were lysed in sample buffer and run on SDS-PAGE gels. WB with anti-KLF1 (Abcam) and anti-actin (Sigma) antibodies was then used to analyze and compare the levels of KLF1 and actin proteins.

### *In vivo* bioluminescence imaging

*Klf1*(K74R) and WT B6 mice were physically restrained and 1×10^5^ B16-F10-luc cells/mouse were intravenously injected into their tail vein. Ten days after melanoma cell inoculation, mice were anesthetized for 5 min and injected intraperitoneally with D-luciferin (300 mg/Kg of body weight). Fifteen minutes after maximum luciferin uptake, the mice were subjected to imaging of the lung and liver regions in an IVIS 50 Bioluminescence imager (Caliper Life Sciences) to determine metastatic burden. The same mice were used the next day as recipients of bone marrow transplantation (BMT) from donor WT or *Klf1*(K74R) mice. Following BMT, bioluminescence imaging was performed on days 0, 10, 17 and 24.

### Bone marrow transplantation (BMT)

BMT followed the standard protocol described in Imado *et al.* (2004)^24^. B6, CD45.1 or CD45.2 donor mice were sacrificed and their femurs were removed. Bone marrow cells were harvested by flashing the femurs with RPM I1640 medium (GIBCO) using a 27-gauge needle and syringe. The cells were then incubated at 37 °C for 30 min in murine complement buffer containing antibodies against B cells, T cells and NK cells, washed twice with PBS, and then subjected to Ficoll-Paque PLUS gradient centrifugation to collect bone marrow mononuclear cells (BMMNCs). BMMNCs (1×10^6^ cells/mouse) from donor mice were injected into the tail veins of recipient B6, CD45.2 or CD45.1 mice that had been exposed to total body γ-irradiation of 10, 5 or 2.5 Gy.

### Calcein-AM cytotoxicity assay of NK cells

1 × 10^6^ target cells (B16-F10 and Hepa1-6) were suspended in 1ml DPBS containing 5 µM Calcein-AM (BioLegend) and incubated at 37℃ for 30 minutes in the dark. The target cells were then washed by RPMI for three times and seeded at 0.3 x 10^5^ cells per well on V-bottom 96-well plate. NK cells from the spleen were sorted by the surface markers, NK1.1^+^, CD3^-^, B220^-^ and NKp46^+^ (Supplementary file 1a), centrifuged at 500 g for 5 minutes, and cultured with IL-15 in RPMI. They were co-cultured with the target cells through different E (Effector): T (Target) ratios (0.5:1, 1:1, 3:1 and 9:1). The plates were centrifuged at 250 g for 1 minute to accelerate the contact of NK cells and target cells and then co-cultured for 4hrs at 37 ℃. Spontaneous release of Calcein-AM was measured in the target cells without NK cells and the maximal release was determined by complete lysis of the target cells in RPMI containing 2% Triton-X 100. After co-culturing, the plates were centrifuged at 120 g for 1 minute and 100 µl/ well of the supernatant was transferred to 96-well assay plate (Corning). The 488/520 nm values were determined by EnSpire (PerkinElmer).

### Bead-based multiplex assay of serum cytokines

Serum samples were obtained via submandibular blood collection and allowed to clot in uncoated tubes for two hours at room temperature. The tubes were centrifuged at 6,000 rpm and the supernatants were collected for cytokine analysis by bead-based multiplex assay (MILLIPLEX MAP Mouse High Sensitivity T Cell Panel, Millipore) following the manufacturer protocol^27^.

### Protein extraction

The cell pellets were resuspended in protein extraction buffer (20 mM HEPES, 0.2% SDS, 1 mM EDTA, 1 mM glycerophosphate, 1 mM Na_3_VO_4_, and 2.5 mM Na_4_P_2_O_7_) with protease inhibitor cocktail (Sigma-Aldrich) and 1 mM phenylmethylsulfonyl fluoride (PMSF). The lysates were further homogenized using a Bioruptor (Diagenode) at 4 °C for 15 min, and then centrifuged at 14,000 × *g* at 4 °C for 20 min. The supernatant was transferred to a new tube before determining protein concentration by means of BCA protein assay (Pierce, Thermo Fisher). Protein aliquots were stored at −30 °C until use.

### In-solution digestion

Protein solutions were first diluted with 50 mM ammonium bicarbonate (ABC) and reduced with 5 mM dithiothreitol (DTT, Merck) at 60 °C for 45 min, followed by cysteine-blocking with 10 mM iodoacetamide (IAM, Sigma) at 25°C for 30 min. The samples were then diluted with 25 mM ABC and digested with sequencing-grade modified porcine trypsin (Promega) at 37 °C for 16 h. The peptides were desalted using a homemade C18 microcolumn (SOURCE 15RPC, GE Healthcare) and stored at −30 °C until use.

### LC-MS/MS analysis

The desalted peptides were diluted in HPLC buffer A (0.1% formic acid in 30% acetonitrile) and loaded onto a homemade SCX column (0.6 × 5 mm, Luna 5 µm SCX 100 Å, Phenomenex). The eluted peptides were then trapped in a reverse-phase column (Zorbax 300SB-C18, 0.3 × 5 mm; Agilent Technologies), and separated on a homemade column (HydroRP 2.5 µm, 75 µm I.D. × 15 cm with a 15 µm tip) using a multi-step gradient of HPLC buffer B (99.9% acetonitrile/0.1% formic acid) for 90 min with a flow rate of 0.3 µl/min. The LC apparatus was coupled to a 2D linear ion trap mass spectrometer (Orbitrap Elite ETD; Thermo Fisher) operated using Xcalibur 2.2 software (Thermo Fisher). Full-scan MS was performed in the Orbitrap over a range of 400 to 2,000 Da and a resolution of 120,000 at m/z 400. Internal calibration was performed using the ion signal of [Si(CH_3_)_2_O]6H^+^ at m/z 536.165365 as lock mass. The 20 data-dependent MS/MS scan events were followed by one MS scan for the 20 most abundant precursor ions in the preview MS scan. The m/z values selected for MS/MS were dynamically excluded for 40 sec with a relative mass window of 10 ppm. The electrospray voltage was set to 2.0 kV, and the temperature of the capillary was set to 200 °C. MS and MS/MS automatic gain control was set to 1,000 ms (full scan) and 200 ms (MS/MS), or to 3 × 106 ions (full scan) and 3,000 ions (MS/MS), for maximum accumulated time or ions, respectively.

### Protein identification

Data analysis was carried out using Proteome Discoverer software (version 1.4, Thermo Fisher Scientific). The MS/MS spectra were searched against the SwissProt database using the Mascot search engine (Matrix Science, version 2.5). For peptide identification, 10 ppm mass tolerance was permitted for intact peptide masses, and 0.5 Da for CID fragment ions with an allowance for two missed cleavages arising from trypsin digestion, oxidized methionine and acetyl (protein N-terminal) as variable modifications, and carbamidomethyl (cysteine) as a static modification. Peptide spectrum matches (PSM) were then filtered based on high confidence and a Mascot search engine ranking of 1 for peptide identification to ensure an overall false discovery rate <0.01. Proteins with single peptide hits were removed from further analysis.

### Gene Set Enrichment Analysis

The absolute abundance of each peptide was calculated from respective peak intensity based on the PSM abundance. The protein abundance of each sample was calculated from the sum of the peptide abundance. The abundance data were then background-corrected and normalized according to variance stabilizing transformation by using the function “normalize_vsn” in the R package *DEP*^70^. Differential expression across groups was determined using the function “test_diff” based on protein-wise linear models combined with empirical Bayes statistics. Significantly differentially-expressed proteins were determined according to a P-value threshold of 0.01 and a fold-change (FC) >1.5. To establish functional pathways enriched across groups, normalized data for each pair of compared groups were used to perform Gene Set Enrichment Analysis (GSEA v4.2.0)^71^ on selected MSigDB gene sets, including Hallmark (H), curated (C2), and immunologic signature (C7) gene sets, by using the default parameters. Normalized enrichment scores (NES) were used to plot a heatmap in the R package *pheatmap* (v1.0.12).

### Statistical analysis

Data are shown as mean ± standard deviation (SD) or standard error of the mean (SEM). Comparisons of data under different experimental conditions were carried out using GraphPad Prism 6.0 software (GraphPad). Each error bar represents SEM unless otherwise indicated. Significant differences in tumor growth on mouse lungs were assessed by Student’s t test. A difference between groups was considered statistically significant when the p value was lower than 0.05.

---

**Supplementary Figure 1-figure supplement 1. Cancer resistance of *Klf1*(K74R) homozygous and eterozygous mice**

(A) Flow chart illustrating the strategy of the pulmonary tumor foci assay. (B) Representative photographs of pulmonary metastatic foci on the lungs of WT, heterozygous *Klf1*(K74R), and homozygous *Klf1*(K74R) male mice in the B6 background two weeks after intravenous injection of B16-F10 melanoma cells (10^5^ cells/ mouse). (C) Statistical comparison of the numbers of pulmonary foci is shown in the three histograms. N=12 (heterozygous), N=7 (WT), and N=6 (homozygous), **, p<0.01. Note that only the numbers of large pulmonary foci (>1mm diameter) were scored.

**Supplementary Figure 2-figure supplement 1.** (A) Transfer of the anti-metastasis capability of 24 month-old male *Klf1*(K74R) mice to 2 month-old male WT mice by BMT with use of 10 Gy γ-irradiation. Statistical comparison of the numbers of pulmonary foci is shown in the two histograms on the right. N=10 (male), *, p<0.05. Note that only the numbers of large pulmonary foci (>1mm diameter) were scored. (B) Survival curves of 2 month-old WT mice receiving BMT from 2 month-old *Klf1*(K74R) or WT donor mice. N=10, p<0.05.

**Supplementary Figure 4-figure supplement 1. <u>Comparison of hematopoietic cell populations between WT and *Klf1*(K74R) mice</u>**

Line graph for comparison of the percentages of peripheral CD3^+^ T cells, B220^+^ B cells, monocytes, granulocytes, and NK cells, respectively, in the peripheral blood of 3 month- old, 12 month-old and 24 month-old WT vs *Klf1*(K74R) male mice. N>5. *, p<0.05.

**Supplementary Figure 4-figure supplement 2. <u>Comparative analysis of *Klf1* gene expression in hematopoietic cells of WT and *Klf1*(K74R) mice</u>**

(A) Comparative Western blotting (WB) and RT-qPCR analysis of *Klf1* expression in leukocytes isolated from peripheral blood of 3 month-old WT and *Klf1* (K74R) mice. The representative WB gel patterns of leukocytes/ WBC are shown in the left panels and the statistical analysis of the RT-qPCR data of *Klf1* mRNA in the leukocytes/ WBC is shown in the right diagram. N=3. (B) KLF1 protein expression in NK cells, CD3^+^ T cells, and B220^+^ B cells isolated from spleen were compared by Western blotting. Extract from mouse erythroleukemia cell line MEL was used as a positive control. The extracts from different types of cells were first standardized by analysis with use of anti-actin antibody (lower panel on the left). They were then analyzed by anti-KLF1. Note that the amount/lane of extracts from NK, T and B cells were all 12 folds higher than the MEL cell extract loaded considering the relatively much higher level of KLF1 in MEL cells. N>4. (C) Comparison of the *Klf1* mRNA levels in CD3^+^ T cells and B220^+^ B cells isolated from the PB of 3 and 24 month-old WT and *Klf1*(K74R) male mice. N=3. *Klf1* mRNA of the MEL cells was also analyzed as the positive control. (D) RT-qPCR was used to compare the levels of *Pdcd* and *Cd274* mRNAs in the peripheral blood of donor WT and *Klf1*(K74R) mice before BMT and the peripheral blood of recipient WT mice after BMT. N=4. *, p<0.05; **, p<0.01.

**Supplementary Figure 4-figure supplement 3. <u>ChIP-qPCR analysis of KLF1-binding on the *Pdcd* promoter of CD3^+^ T cells from WT and *Klf1*(K74R) mice</u>**

The map of the *Pdcd* promoter region is shown on top, with the CACCC boxes (Box1, Box2, Box3), CCAAT boxes (C1), and the transcription start site (TSS, +1) indicated. The histogram below shows the relative signals from ChIP-qPCR analysis of the regions a, b, and c centered around the Box1, Box2, and Box3, respectively. Note that KLF1 binds to Box3 of *Pdcd* promoter in CD3^+^ T cells from either WT or *Klf1*(K74R) mice. ChIP-qPCR analysis of KLF1-binding in the *ýmaj* globin gene promoter in MEL cells was carried out as a positive control (data not shown). N=2, * p< 0.05.

**Supplementary Figure 4-figure supplement 4. <u>Comparison of the serum levels of different cytokines in WT and *Klf1*(K74R) mice</u>**

Bead-based multiplex analysis was used to measure the serum levels of IL-1β, IL-2, IL-4, IL-6, IL-10, IL-12p70, TNF-α and IFN-γ in the sera of the male mice of the ages of 3 months and 24 months. N>3. *, p<0.05; **, p<0.01; ***, p<0.001.

**Supplementary file 1a. <u>Cell surface markers of different hematopoietic blood cells</u>**

**Supplementary file 1b. <u>DNA primers used in ChIP-qPCR experiments</u>**

**Figure 4-figure supplement 2-source data 1 Original file for the Western blot analysis in Figure 4-figure supplement 2A (anti-KLF1, anti-β-Actin).**

**Figure 4-figure supplement 2-source data 2 Original file for the Western blot analysis in Figure 4-figure supplement 2B (anti-KLF1, anti-β-Actin).**

## References

1. Aman Y, Schmauck-Medina T, Hansen M, et al. Autophagy in healthy aging and disease. Nat Aging. Aug 2021;1(8):634–650. doi:10.1038/s43587-021-00098-4

2. López-Otín C, Blasco MA, Partridge L, Serrano M, Kroemer G. The hallmarks of aging. Cell. Jun 6 2013;153(6):1194–217. doi:10.1016/j.cell.2013.05.039

3. Bashor CJ, Hilton IB, Bandukwala H, Smith DM, Veiseh O. Engineering the next generation of cell-based therapeutics. Nat Rev Drug Discov. Sep 2022;21(9):655–675. doi:10.1038/s41573-022-00476-6

4. Fontana L, Partridge L. Promoting health and longevity through diet: from model organisms to humans. Cell. Mar 26 2015;161(1):106–118. doi:10.1016/j.cell.2015.02.020

5. Longo VD, Anderson RM. Nutrition, longevity and disease: From molecular mechanisms to interventions. Cell. Apr 28 2022;185(9):1455–1470. doi:10.1016/j.cell.2022.04.002

6. Mitchell E, Spencer Chapman M, Williams N, et al. Clonal dynamics of haematopoiesis across the human lifespan. Nature. Jun 2022;606(7913):343–350. doi:10.1038/s41586-022-04786-y

7. Pálovics R, Keller A, Schaum N, et al. Molecular hallmarks of heterochronic parabiosis at single-cell resolution. Nature. Mar 2022;603(7900):309–314. doi:10.1038/s41586-022-04461-2

8. Seita J, Weissman IL. Hematopoietic stem cell: self-renewal versus differentiation. Wiley Interdiscip Rev Syst Biol Med. Nov-Dec 2010;2(6):640–53. doi:10.1002/wsbm.86

9. Montazersaheb S, Ehsani A, Fathi E, Farahzadi R. Cellular and Molecular Mechanisms Involved in Hematopoietic Stem Cell Aging as a Clinical Prospect. Oxid Med Cell Longev. 2022;2022:2713483. doi:10.1155/2022/2713483

10. Frontelo P, Manwani D, Galdass M, et al. Novel role for EKLF in megakaryocyte lineage commitment. Blood. Dec 1 2007;110(12):3871–80. doi:10.1182/blood-2007-03-082065

11. Nishizawa M, Chonabayashi K, Nomura M, et al. Epigenetic Variation between Human Induced Pluripotent Stem Cell Lines Is an Indicator of Differentiation Capacity. Cell Stem Cell. Sep 1 2016;19(3):341–54. doi:10.1016/j.stem.2016.06.019

12. Teruya S, Okamura T, Komai T, et al. Egr2-independent, Klf1-mediated induction of PD-L1 in CD4(+) T cells. Sci Rep. May 4 2018;8(1):7021. doi:10.1038/s41598-018-25302-1

13. Perkins A, Xu X, Higgs DR, et al. Krüppeling erythropoiesis: an unexpected broad spectrum of human red blood cell disorders due to KLF1 variants. Blood. Apr 14 2016;127(15):1856–62. doi:10.1182/blood-2016-01-694331

14. Neuwirtova R, Fuchs O, Holicka M, et al. Transcription factors Fli1 and EKLF in the differentiation of megakaryocytic and erythroid progenitor in 5q-syndrome and in Diamond-Blackfan anemia. Ann Hematol. Jan 2013;92(1):11–8. doi:10.1007/s00277-012-1568-1

15. Luo Q, Ma X, Wahl SM, Bieker JJ, Crossley M, Montaner LJ. Activation and repression of interleukin-12 p40 transcription by erythroid Kruppel-like factor in macrophages. J Biol Chem. Apr 30 2004;279(18):18451–6. doi:10.1074/jbc.M400320200

16. Hung CH, Wang KY, Liou YH, et al. Negative Regulation of the Differentiation of Flk2(-) CD34(-) LSK Hematopoietic Stem Cells by EKLF/KLF1. Int J Mol Sci. Nov 10 2020;21(22)doi:10.3390/ijms21228448

17. Siatecka M, Bieker JJ. The multifunctional role of EKLF/KLF1 during erythropoiesis. Blood. Aug 25 2011;118(8):2044–54. doi:10.1182/blood-2011-03-331371

18. Pilon AM, Ajay SS, Kumar SA, et al. Genome-wide ChIP-Seq reveals a dramatic shift in the binding of the transcription factor erythroid Kruppel-like factor during erythrocyte differentiation. Blood. Oct 27 2011;118(17):e139–48. doi:10.1182/blood-2011-05-355107

19. Tallack MR, Magor GW, Dartigues B, et al. Novel roles for KLF1 in erythropoiesis revealed by mRNA-seq. Genome Res. Dec 2012;22(12):2385–98. doi:10.1101/gr.135707.111

20. Shyu YC, Lee TL, Chen X, et al. Tight regulation of a timed nuclear import wave of EKLF by PKCθ and FOE during Pro-E to Baso-E transition. Dev Cell. Feb 24 2014;28(4):409–22. doi:10.1016/j.devcel.2014.01.007

21. Shyu YC, Liao PC, Huang TS, et al. Genetic Disruption of KLF1 K74 SUMOylation in Hematopoietic System Promotes Healthy Longevity in Mice. Adv Sci (Weinh). Sep 2022;9(25):e2201409. doi:10.1002/advs.202201409

22. Vaseghi G, Dana N, Ghasemi A, Abediny R, Laher I, Javanmard SH. Morphine promotes migration and lung metastasis of mouse melanoma cells. Braz J Anesthesiol. Jul-Aug 2023;73(4):441–445. doi:10.1016/j.bjane.2021.10.019

23. Díaz-García VM, Guerrero S, Díaz-Valdivia N, et al. Biomimetic quantum dot-labeled B16F10 murine melanoma cells as a tool to monitor early steps of lung metastasis by in vivo imaging. Int J Nanomedicine. 2018;13:6391–6412. doi:10.2147/ijn.S165565

24. Imado T, Iwasaki T, Kataoka Y, et al. Hepatocyte growth factor preserves graft-versus-leukemia effect and T-cell reconstitution after marrow transplantation. Blood. Sep 1 2004;104(5):1542–9. doi:10.1182/blood-2003-12-4309

25. Iwai Y, Hamanishi J, Chamoto K, Honjo T. Cancer immunotherapies targeting the PD-1 signaling pathway. J Biomed Sci. Apr 4 2017;24(1):26. doi:10.1186/s12929-017-0329-9

26. Jiang X, Wang J, Deng X, et al. Role of the tumor microenvironment in PD-L1/PD-1-mediated tumor immune escape. Mol Cancer. Jan 15 2019;18(1):10. doi:10.1186/s12943-018-0928-4

27. Aira C, Ruiz T, Dixon L, Blome S, Rueda P, Sastre P. Bead-Based Multiplex Assay for the Simultaneous Detection of Antibodies to African Swine Fever Virus and Classical Swine Fever Virus. Front Vet Sci. 2019;6:306. doi:10.3389/fvets.2019.00306

28. Menees KB, Earls RH, Chung J, et al. Sex- and age-dependent alterations of splenic immune cell profile and NK cell phenotypes and function in C57BL/6J mice. Immun Ageing. Jan 8 2021;18(1):3. doi:10.1186/s12979-021-00214-3

29. Ul-Haq Z, Naz S, Mesaik MA. Interleukin-4 receptor signaling and its binding mechanism: A therapeutic insight from inhibitors tool box. Cytokine Growth Factor Rev. Dec 2016;32:3–15. doi:10.1016/j.cytogfr.2016.04.002

30. Boccardi V, Westman E, Pelini L, et al. Differential Associations of IL-4 With Hippocampal Subfields in Mild Cognitive Impairment and Alzheimer’s Disease. Front Aging Neurosci. 2018;10:439. doi:10.3389/fnagi.2018.00439

31. Rea IM, Gibson DS, McGilligan V, McNerlan SE, Alexander HD, Ross OA. Age and Age-Related Diseases: Role of Inflammation Triggers and Cytokines. Front Immunol. 2018;9:586. doi:10.3389/fimmu.2018.00586

32. Fabian DK, Fuentealba M, Dönertaş HM, Partridge L, Thornton JM. Functional conservation in genes and pathways linking ageing and immunity. Immun Ageing. May 14 2021;18(1):23. doi:10.1186/s12979-021-00232-1

33. Yeo EJ. Hypoxia and aging. Exp Mol Med. Jun 20 2019;51(6):1-15. doi:10.1038/s12276-019-0233-3

34. Zhu X, Chen Z, Shen W, et al. Inflammation, epigenetics, and metabolism converge to cell senescence and ageing: the regulation and intervention. Signal Transduct Target Ther. Jun 28 2021;6(1):245. doi:10.1038/s41392-021-00646-9

35. Hamarsheh S, Groß O, Brummer T, Zeiser R. Immune modulatory effects of oncogenic KRAS in cancer. Nat Commun. Oct 28 2020;11(1):5439. doi:10.1038/s41467-020-19288-6

36. Ni L, Lu J. Interferon gamma in cancer immunotherapy. Cancer Med. Sep 2018;7(9):4509–4516. doi:10.1002/cam4.1700

37. Principe DR, Doll JA, Bauer J, et al. TGF-β: duality of function between tumor prevention and carcinogenesis. J Natl Cancer Inst. Feb 2014;106(2):djt369. doi:10.1093/jnci/djt369

38. Zitvogel L, Galluzzi L, Kepp O, Smyth MJ, Kroemer G. Type I interferons in anticancer immunity. Nat Rev Immunol. Jul 2015;15(7):405–14. doi:10.1038/nri3845

39. Zhao G, Zhu G, Huang Y, et al. IL-6 mediates the signal pathway of JAK-STAT3-VEGF-C promoting growth, invasion and lymphangiogenesis in gastric cancer. Oncol Rep. Mar 2016;35(3):1787–95. doi:10.3892/or.2016.4544

40. Roncagalli R, Taylor JE, Zhang S, et al. Negative regulation of natural killer cell function by EAT-2, a SAP-related adaptor. Nat Immunol. Oct 2005;6(10):1002–10. doi:10.1038/ni1242

41. Yu Y. The Function of NK Cells in Tumor Metastasis and NK Cell-Based Immunotherapy. Cancers (Basel). Apr 16 2023;15(8)doi:10.3390/cancers15082323

42. Ferreira FM, Palle P, Vom Berg J, Prajwal P, Laman JD, Buch T. Bone marrow chimeras-a vital tool in basic and translational research. J Mol Med (Berl). Jul 2019;97(7):889–896. doi:10.1007/s00109-019-01783-z

43. Bin-Jumah MN, Nadeem MS, Gilani SJ, et al. Genes and Longevity of Lifespan. Int J Mol Sci. Jan 28 2022;23(3)doi:10.3390/ijms23031499

44. Hofmann JW, Zhao X, De Cecco M, et al. Reduced expression of MYC increases longevity and enhances healthspan. Cell. Jan 29 2015;160(3):477–88. doi:10.1016/j.cell.2014.12.016

45. Ocampo A, Reddy P, Martinez-Redondo P, et al. In Vivo Amelioration of Age-Associated Hallmarks by Partial Reprogramming. Cell. Dec 15 2016;167(7):1719–1733.e12. doi:10.1016/j.cell.2016.11.052

46. Varady KA, Cienfuegos S, Ezpeleta M, Gabel K. Clinical application of intermittent fasting for weight loss: progress and future directions. Nat Rev Endocrinol. May 2022;18(5):309–321. doi:10.1038/s41574-022-00638-x

47. Hifumi T, Yamamoto A, Ato M, et al. Clinical Serum Therapy: Benefits, Cautions, and Potential Applications. Keio J Med. Dec 25 2017;66(4):57–64. doi:10.2302/kjm.2016-0017-IR

48. Morgan RA, Gray D, Lomova A, Kohn DB. Hematopoietic Stem Cell Gene Therapy: Progress and Lessons Learned. Cell Stem Cell. Nov 2 2017;21(5):574–590. doi:10.1016/j.stem.2017.10.010

49. Giannaccare G, Carnevali A, Senni C, Logozzo L, Scorcia V. Umbilical Cord Blood and Serum for the Treatment of Ocular Diseases: A Comprehensive Review. Ophthalmol Ther. Jun 2020;9(2):235–248. doi:10.1007/s40123-020-00239-9

50. Wu SY, Fu T, Jiang YZ, Shao ZM. Natural killer cells in cancer biology and therapy. Mol Cancer. Aug 6 2020;19(1):120. doi:10.1186/s12943-020-01238-x

51. Daver N, Wei AH, Pollyea DA, Fathi AT, Vyas P, DiNardo CD. New directions for emerging therapies in acute myeloid leukemia: the next chapter. Blood Cancer J. Oct 30 2020;10(10):107. doi:10.1038/s41408-020-00376-1

52. Mora J. Autologous Stem-Cell Transplantation for High-Risk Neuroblastoma: Historical and Critical Review. Cancers (Basel). May 24 2022;14(11)doi:10.3390/cancers14112572

53. Conboy IM, Conboy MJ, Wagers AJ, Girma ER, Weissman IL, Rando TA. Rejuvenation of aged progenitor cells by exposure to a young systemic environment. Nature. Feb 17 2005;433(7027):760-4. doi:10.1038/nature03260

54. Das MM, Godoy M, Chen S, et al. Young bone marrow transplantation preserves learning and memory in old mice. Commun Biol. 2019;2:73. doi:10.1038/s42003-019-0298-5

55. Goodell MA, Rando TA. Stem cells and healthy aging. Science. Dec 4 2015;350(6265):1199-204. doi:10.1126/science.aab3388

56. Kang S, Moser VA, Svendsen CN, Goodridge HS. Rejuvenating the blood and bone marrow to slow aging-associated cognitive decline and Alzheimer’s disease. Commun Biol. Feb 13 2020;3(1):69. doi:10.1038/s42003-020-0797-4

57. Castellano JM, Mosher KI, Abbey RJ, et al. Human umbilical cord plasma proteins revitalize hippocampal function in aged mice. Nature. Apr 27 2017;544(7651):488-492. doi:10.1038/nature22067

58. Mehdipour M, Skinner C, Wong N, et al. Rejuvenation of three germ layers tissues by exchanging old blood plasma with saline-albumin. Aging (Albany NY). May 30 2020;12(10):8790–8819. doi:10.18632/aging.103418

59. Siatecka M, Xue L, Bieker JJ. Sumoylation of EKLF promotes transcriptional repression and is involved in inhibition of megakaryopoiesis. Mol Cell Biol. Dec 2007;27(24):8547–60. doi:10.1128/mcb.00589-07

60. Aunan JR, Cho WC, Søreide K. The Biology of Aging and Cancer: A Brief Overview of Shared and Divergent Molecular Hallmarks. Aging Dis. Oct 2017;8(5):628–642. doi:10.14336/ad.2017.0103

61. Lages CS, Lewkowich I, Sproles A, Wills-Karp M, Chougnet C. Partial restoration of T-cell function in aged mice by in vitro blockade of the PD-1/PD-L1 pathway. Aging Cell. Oct 2010;9(5):785–98. doi:10.1111/j.1474-9726.2010.00611.x

62. Schaum N, Lehallier B, Hahn O, et al. Ageing hallmarks exhibit organ-specific temporal signatures. Nature. Jul 2020;583(7817):596-602. doi:10.1038/s41586-020-2499-y

63. Aira C, Ruiz T, Dixon L, Blome S, Rueda P, Sastre P. Bead-Based Multiplex Assay for the Simultaneous Detection of Antibodies to African Swine Fever Virus and Classical Swine Fever Virus. Original Research. Frontiers in Veterinary Science. 2019-September-13 2019;6 doi:10.3389/fvets.2019.00306

64. Zhou J, Zhu Z, Bai C, Sun H, Wang X. Proteomic profiling of lymphocytes in autoimmunity, inflammation and cancer. J Transl Med. Jan 7 2014;12:6. doi:10.1186/1479-5876-12-6

65. Zhang D, Chen G, Manwani D, et al. Neutrophil ageing is regulated by the microbiome. Nature. Sep 24 2015;525(7570):528-32. doi:10.1038/nature15367

66. Narasimhan PB, Eggert T, Zhu YP, et al. Patrolling Monocytes Control NK Cell Expression of Activating and Stimulatory Receptors to Curtail Lung Metastases. J Immunol. Jan 1 2020;204(1):192–198. doi:10.4049/jimmunol.1900998

67. Han Z, Yang D, Trivett A, Oppenheim JJ. Therapeutic vaccine to cure large mouse hepatocellular carcinomas. Oncotarget. Aug 8 2017;8(32):52061–52071. doi:10.18632/oncotarget.19367

68. Daftari P, Gavva NR, Shen CK. Distinction between AP1 and NF-E2 factor-binding at specific chromatin regions in mammalian cells. Oncogene. Sep 23 1999;18(39):5482–6. doi:10.1038/sj.onc.1202916

69. Green MR, Sambrook J. Molecular Cloning: A Laboratory Manual. Cold Spring Harbor Laboratory Press; 2012.

70. Zhang X, Smits AH, van Tilburg GB, Ovaa H, Huber W, Vermeulen M. Proteome-wide identification of ubiquitin interactions using UbIA-MS. Nat Protoc. Mar 2018;13(3):530–550. doi:10.1038/nprot.2017.147

71. Subramanian A, Tamayo P, Mootha VK, et al. Gene set enrichment analysis: a knowledge-based approach for interpreting genome-wide expression profiles. Proc Natl Acad Sci U S A. Oct 25 2005;102(43):15545–50. doi:10.1073/pnas.0506580102

