## Supplementary Figures and Files for "Long-term Hematopoietic Transfer of the Anti-Cancer and Lifespan-Extending Capabilities of A Genetically Engineered Blood System by Transplantation of Bone Marrow Mononuclear Cells"

### Figure 1-figure supplement 1

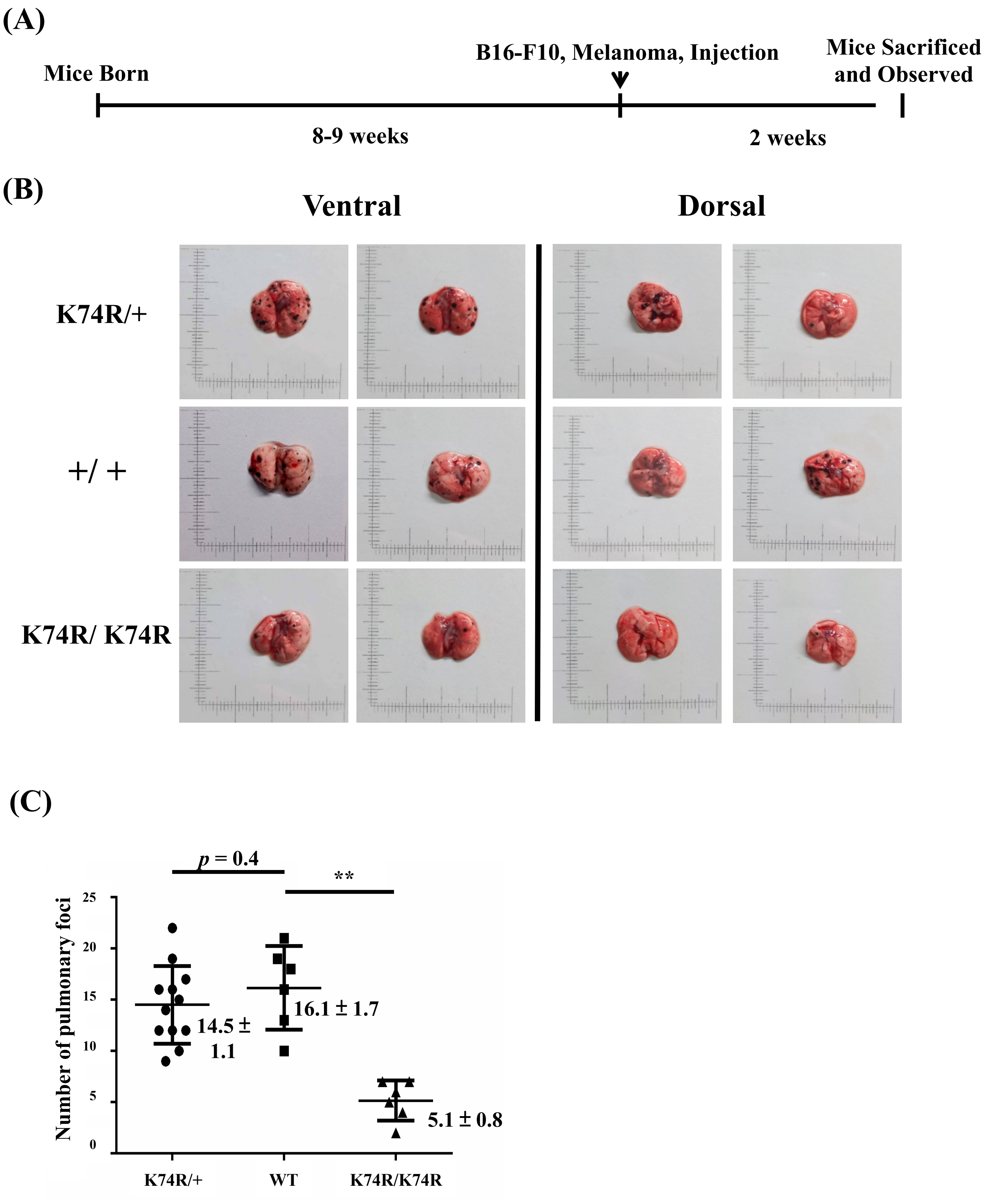

(A)

### Figure 2-figure supplement 1

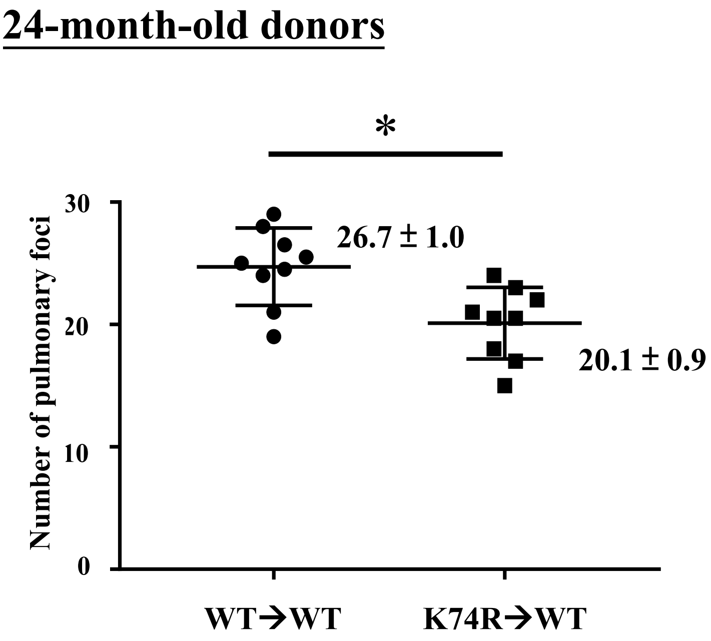


(B)

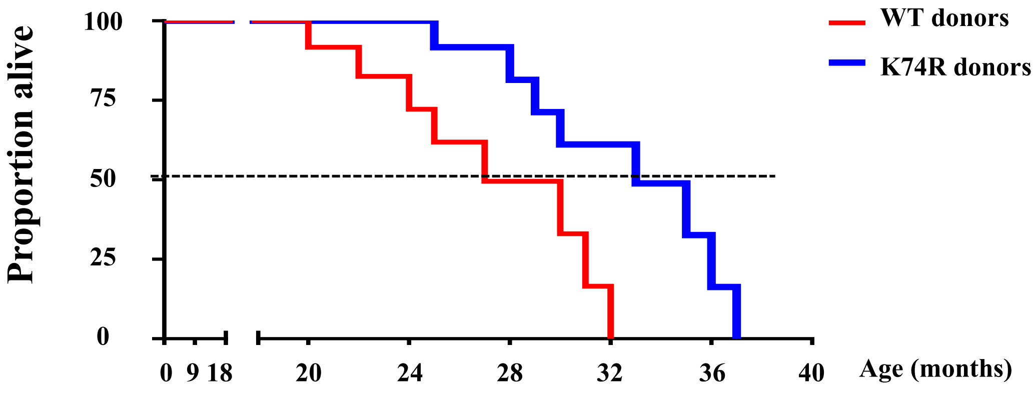


**Life span (Months)**

| | Mediam | Mean | Oldest 10 % | Youngest 10 % | Maximum | Minimum | N |
|---|---|---|---|---|---|---|---|
| WT | 25.9 | 26.5 ± 1.8 | 31.2 ± 1.6 | 20.6 ± 1.9 | 31.3 | 20.5 | 10 |
| K74R | 30.8 | 31.9 ± 1.1 | 35.8 ± 1.9 | 24.7 ± 1.4 | 37.1 | 24.5 | 10 |

### Figure 4-figure supplement 1

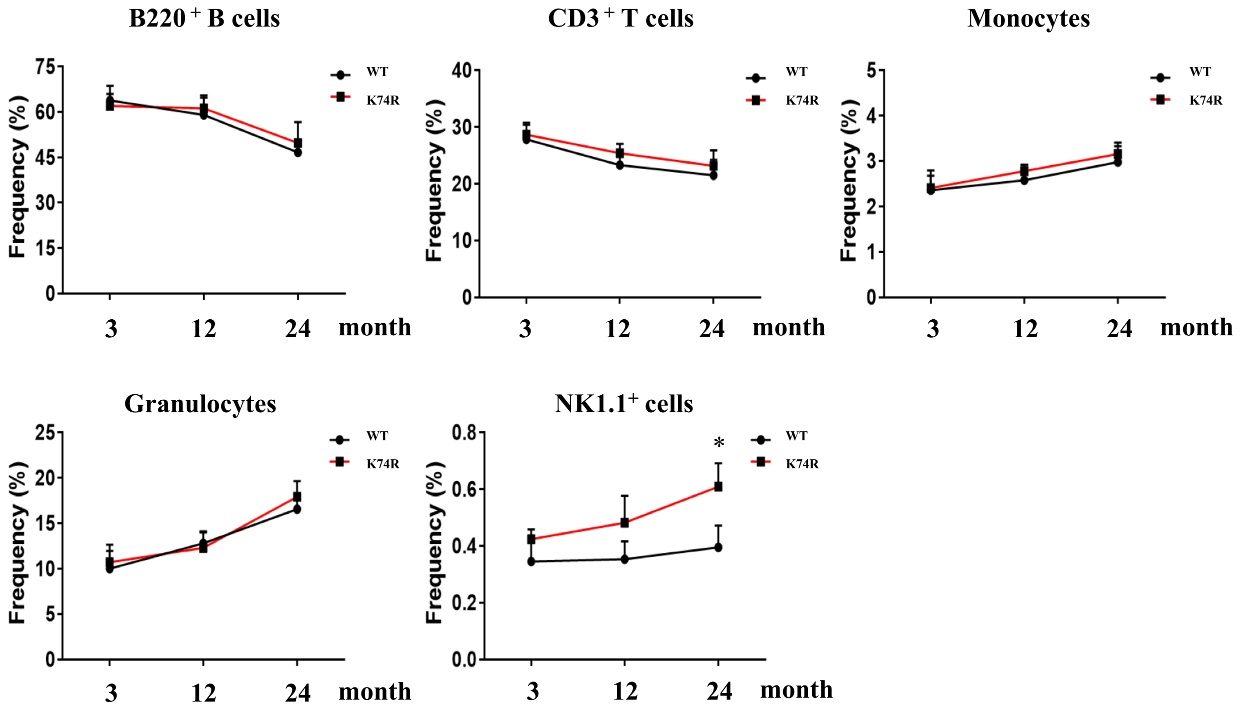

### Figure 4-figure supplement 2

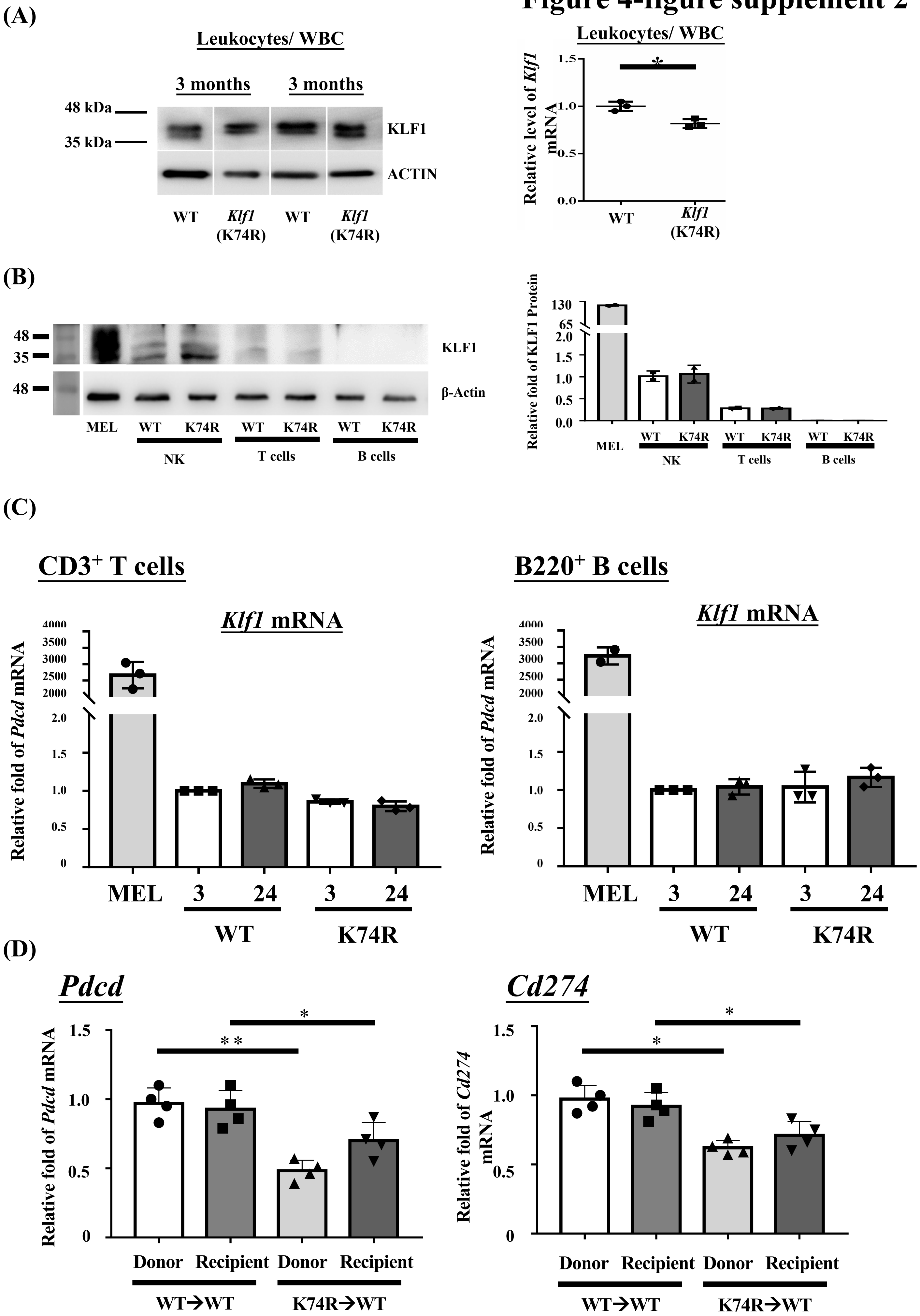

**Figure 4-figure supplement 3**

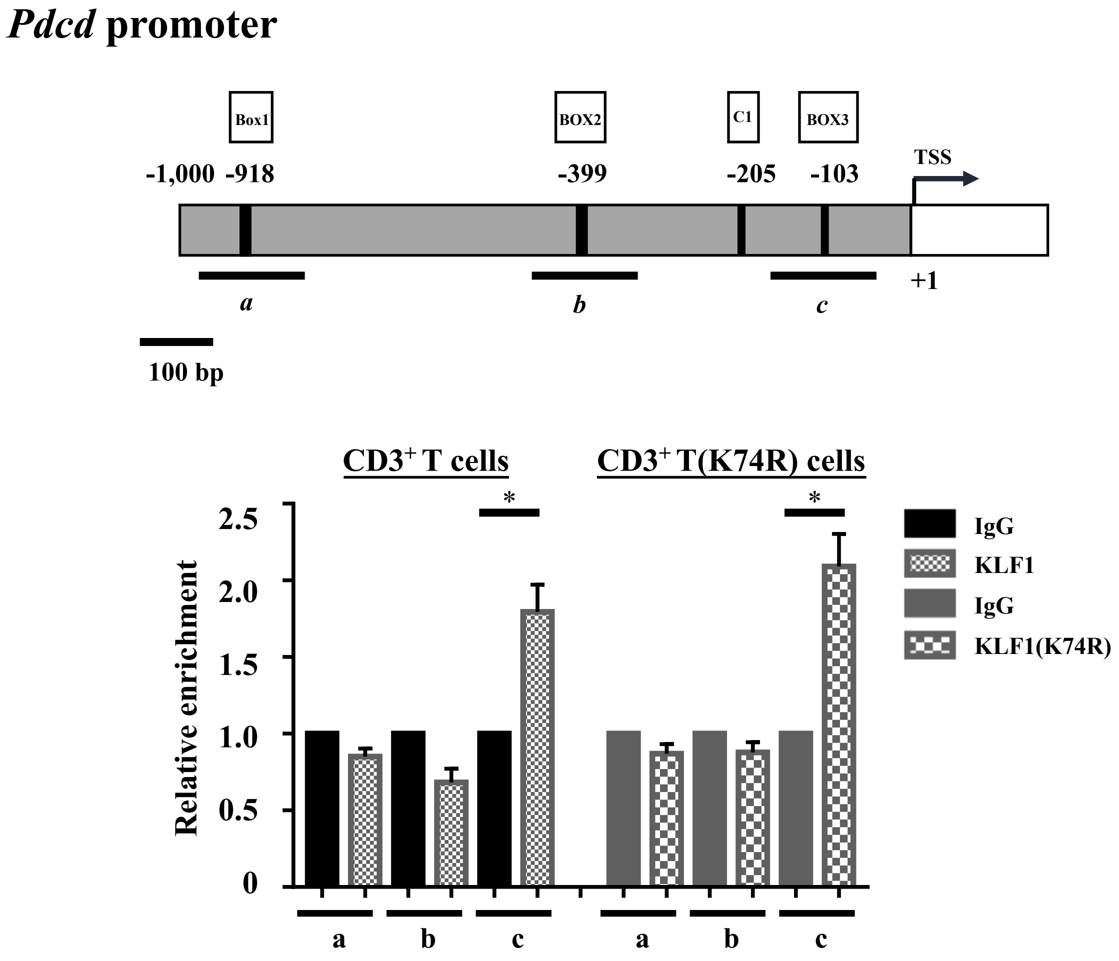

### Figure 4-figure supplement 4

#### Serum

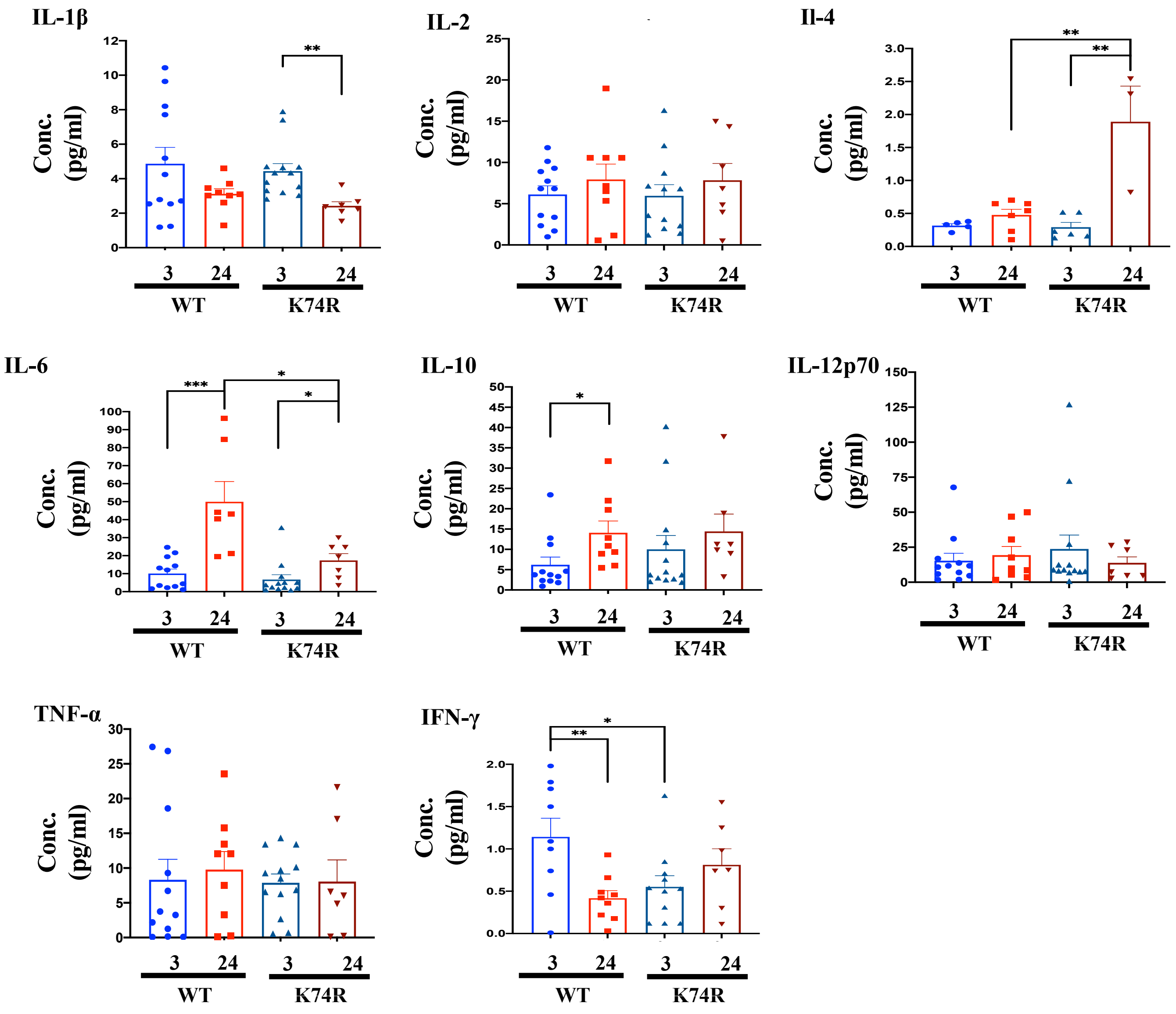

**Supplementary file 1a**

| Marker | Cell types |
| --- | --- |
| $CD45R^{+}/CD3e^{-}$ | B cells |
| $CD45R^{-}/CD3e^{+}$ | T cells |
| $CD45R^{-}/CD3e^{-}/NK1.1^{+}/NKp46^{+}$ | NK cells |
| $CD11b^{+}/Gr\text{-}1^{-}$ | Monocytes |
| $CD11b^{+}/Gr\text{-}1^{+}$ | Granulocytes |

**Supplementary file 1b**

| ChIP-qPCR primers | Sequence 5’ → 3’ |
|---|---|
| Actb+668L | 5’-GAC TAC CTC ATG AAG ATC CTG AC-3’ |
| Actb+817 | 5’-CAG GGA AGA AGA GGA TGC GGC C-3’ |
| βmajor-6411 | 5’-GTT CTC TGC ACA GAT AAG GAC AAA C-3’ |
| βmajor-6623 | 5’-CTG ATC CTA CCT CAC CTT ATA TGC-3’ |
| Pd-1-d-F | 5’-GAC CTA GAA ATT GAG TCT AC-3’ |
| Pd-1-d-R | 5’-CTC TTA AGG CTT TTC TTC CTT TC-3’ |
| Pd-1 Box1-F | 5’-GTA CCA AAG CCA GGC CTC GAC-3’ |
| Pd-1 Box1-R | 5’-CCT CCT GTG GGT AGG TTT GG-3’ |
| Pd-1 Box2-F | 5’-CTC CCC CAC CTC TAG TTG C-3’ |
| Pd-1 Box2-R | 5’-GGC AGA GTT GTC TGT AGC G-3’ |
